# The evolution of non-seasonal breeding in primates

**DOI:** 10.64898/2026.05.19.725975

**Authors:** Lugdiwine Burtschell, Lucie Thel, Jules Dezeure, Dieter Lukas, Bernard Godelle, Elise Huchard

## Abstract

Reproductive seasonality offsets the energetic costs of reproduction by synchronizing births with peak resources and is traditionally expected to increase with latitude and environmental seasonality. However, life-history and behavioural strategies may also buffer energetic shortages and reduce dependency on environmental cycles. Here, we propose and test an integrative framework integrating climatic, life-history and behavioural factors using high-resolution measures of reproductive seasonality for 132 wild primate populations from 94 species. As expected, reproductive seasonality declines at lower latitudes, in less seasonal and in less predictable environments, even after controlling for productivity. It also decreases in species that spread reproductive costs by extending developmental periods, or when a higher infant mortality urges females to resume fertility shortly after loss. Unexpectedly, reproductive seasonality increases with environmental productivity and is not reduced by cognitive (foraging innovations), social (allomaternal care), or ecological (dietary breadth) buffering. Broader diets even enhance seasonality. These findings suggest that reproductive seasonality emerges from opportunity more than from constraints in productive environments, where females exploit abundant resources to accelerate their reproductive pace. Together, our results shed light on the diverse selective pressures shaping primate reproductive seasonality, including climate, life-history pace, and infanticide risk, and help to explain why humans reproduce year-round.

## Introduction

### Background

Reproduction is energetically costly, especially in mammals where females support offspring growth over prolonged periods of time, through gestation and lactation (Speakman, 2008). To offset these energy expenses, a widespread strategy consists in reproducing seasonally (Baker, 1938; Bronson, 1985). In most environments, food availability fluctuates predictably across seasons and species synchronize the most energetically demanding phases of reproduction with peak resource abundance (Thomas et al., 2001; Brockman and van Schaik, 2005). However, reproducing seasonally also entails costs, notably the introduction of reproductive gaps, during which females must wait for the next breeding season to reproduce. Such delays can be particularly costly following the loss of a litter or offspring. Various theories have therefore proposed how mammals might mitigate these trade-offs, but as yet no coherent framework exists or has been tested.

A wide variety of reproductive seasonality patterns is observed among mammals (e.g. Heldstab et al., 2018). For example, 80% of births in mountain goats (*Oreamnos americanus*) occur within two weeks (Côté and Festa-Bianchet, 2001) whereas births are distributed throughout the year in mountain gorillas (*Gorilla gorilla beringei*; Watts, 1998). One of the most robust predictors of this variation is latitude, with species living further from the equator being more seasonal than tropical ones (English et al., 2012; Zerbe et al., 2012; Heldstab et al., 2018; Heldstab, 2021a, 2021b). This relationship has been interpreted as reflecting the higher degree of environmental seasonality observed at higher latitudes, leading to greater differences in food availability between seasons as well as to food peaks that are narrower in time, therefore increasing the benefits of seasonal reproduction.

However, some of this variation is unexplained by latitude, as seasonal breeders are commonly encountered in tropical or equatorial environments, as well as non-seasonal breeders in temperate habitats. For example, the sympatric ungulates from the Serengeti National Park (latitude around 2°S) can reproduce both seasonally and non-seasonally (Sinclair et al., 2000), and coypus (*Myocastor coypus*) do not show any seasonal pattern in their birth distribution, despite living in habitats of relatively high latitudes (around 33°S) with a marked seasonality in rainfall and temperature (Courtalon et al., 2015). Overall, variation in the intensity of reproductive seasonality across mammals cannot be explained by climatic seasonality alone and instead reflects additional determinants that alter the cost-benefit balance of seasonal reproduction.

Several studies have investigated the nature of such additional determinants in multiple taxonomic groups relatively recently, including ungulates (English et al., 2012), ruminants (Zerbe et al., 2012), carnivores (Heldstab et al., 2018), lagomorphs (Heldstab, 2021a), rodents (Heldstab, 2021b) and primates (Heldstab et al., 2021). The range of parameters tested is wide, covering mostly life-history traits (e.g. gestation length, body mass, litter size, age at sexual maturity) but also environmental (e.g. interannual variability, mean precipitations, altitude), ecological (e.g. diet), or social parameters (e.g. gregariousness, mating system). However, such studies often remain relatively descriptive in their design, so that we still lack a theoretical framework that could help us to gain predictive power. In addition, most of these studies use data from captive animals who often live far from their geographical range, and whose reproductive phenology can thus be deeply altered (e.g. the dorcas gazelle (*Gazella dorcas*) displays less seasonality in captivity than in the wild, Zerbe et al., 2012), and/or use low-resolution and categorical measures of reproductive seasonality in the wild, which drastically limit analytical possibilities. Here we propose a general framework to explain the evolution of (non-)seasonal breeding that we assess and refine using a dataset with high resolution measures of reproductive seasonality in wild primates.

### Theoretical framework

Some of our framework proposed here has been inspired from anthropological contributions aimed at understanding the energetic challenges linked with maintaining a large brain (van Woerden, 2011), adjusted and extended to understand the evolution of reproductive seasonality. From an energetic perspective, strictly seasonal breeding—defined by a complete absence of births during part of the year—may be favoured when, at some point of the annual cycle, energy availability falls below the threshold required to sustain the most energetically demanding phase of reproduction. In contrast, non-seasonal breeding may evolve when energy availability consistently exceeds the maximum energetic demands of reproduction throughout the year. Under such conditions, reproduction can be sustained year-round, a situation that may be achieved through several, non-mutually exclusive mechanisms.

First, a higher environmental productivity increases the overall energy available at all times, therefore allowing reproduction for an extended proportion of the annual cycle or even year-round, while the level of reproductive costs remains unchanged (Fig. 1B). Second, and symmetrically, the maximal amount of energy needed for reproduction could be lowered, without modifying the overall energy availability, therefore allowing for energy needs to be met for an extended proportion of the annual cycle, or even throughout the year (Fig. 1C). Specifically, life history strategies consisting in slowing down the reproductive pace may allow species to spread reproductive effort over extended periods (e.g. van Noordwijk et al., 2013), in a way that lowers their daily reproductive costs. On the contrary, species with fast life histories and condensed maternal investment are more likely to exploit resource bursts and thus reproduce seasonally. Lastly, some behavioural strategies could contribute to buffer the seasonality of the environment, by increasing the amount of energy extracted at times of food shortage (Fig. 1D). In such cases, the seasonality of the environment would remain unchanged, but the seasonality experienced by a species would be reduced. “Cognitive buffers” could be a first example of buffering of the environmental seasonality. Enhanced cognitive abilities would enable species to increase their foraging efficiency, for example through the development of foraging innovations (Reader et al., 2011), that would provide them with extra energetic inputs during the lean season. “Ecological buffers” could play a similar role: behavioural flexibility in diet, recourse to fall-back foods during the lean period when usual food items are unavailable (Altmann, 1998), and overall, a wider diet breadth (Kissling et al., 2014) could reduce their vulnerability to starvation. Lastly, “social buffers” covering all forms of allomaternal care, such as cooperative and communal breeding, but particularly through allomaternal provisioning, could also buffer the costs of reproducing during the lean season, by alleviating the maternal costs of reproduction (Lukas and Clutton-Brock, 2017; Groenewoud and Clutton-Brock, 2021). Overall, a high environmental productivity (Fig. 1B), the spread of reproductive effort (Fig. 1C) or the use of cognitive, ecological or social buffers to overcome food shortage (Fig. 1D), could all contribute to shorten or cancel the time period when the energy needs of reproduction cannot be met. In such cases, the benefits of seasonal breeding would be limited or null and the cost-benefit balance would tilt towards non-seasonal reproduction.

**Figure 1:**
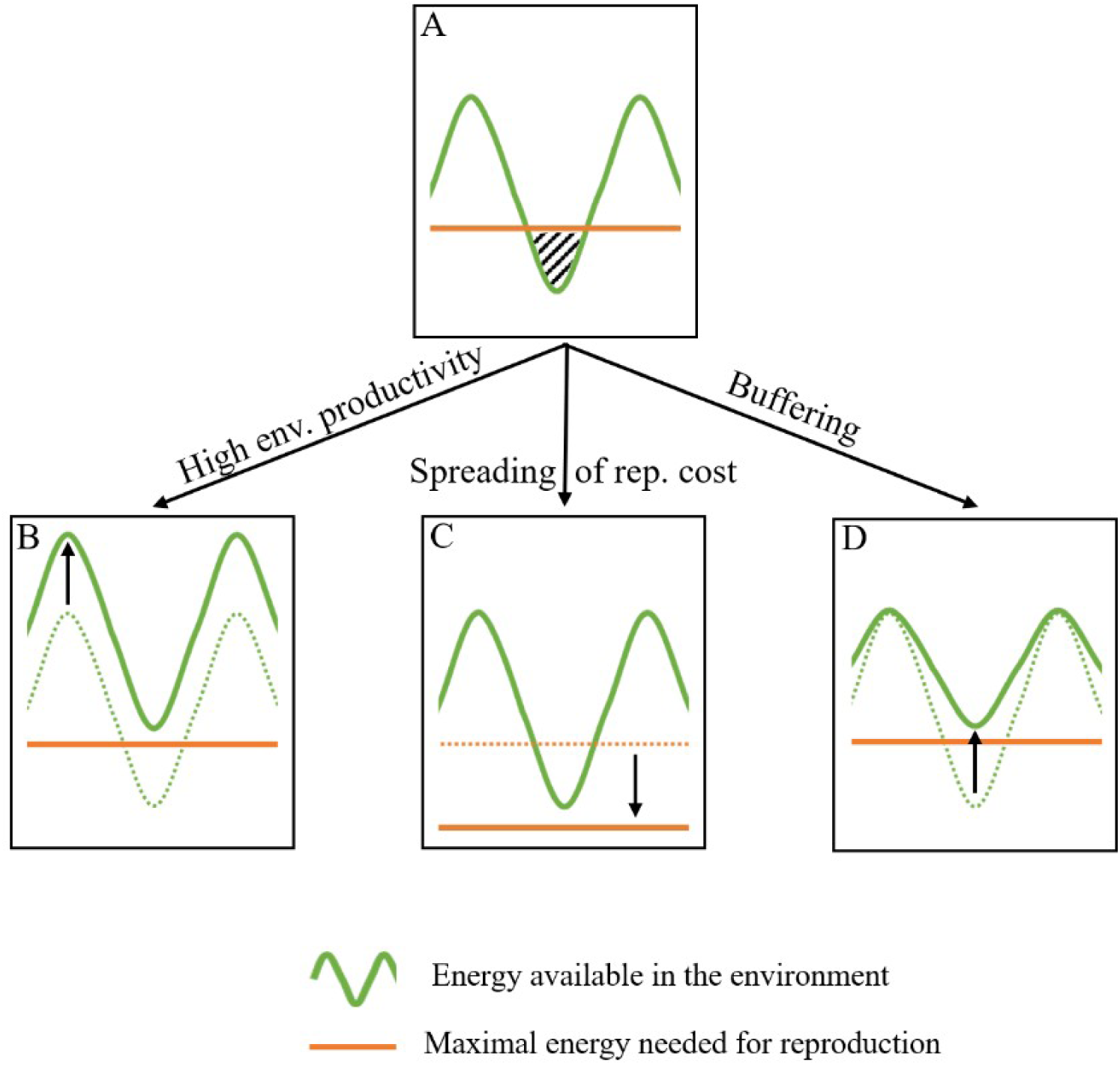
Theoretical representation of a case where seasonal breeding would be favoured (A), with a time period where the energy available in the environment goes below the maximal amount of energy needed for reproduction (hatched area). Strict non-seasonal breeding, where reproduction can happen at all times, may evolve if this maximal amount is always above the available energy. This can be achieved through a higher environmental productivity (B), a spreading of reproductive costs (C) or by behavioural mechanisms buffering energy shortage (D), for example through cognitive, ecological or social buffers. Figure adapted from van Woerden (2011).

In addition to increasing the energy available or decreasing energetic needs, another environmental characteristic, namely environmental unpredictability, could limit the benefits of seasonal reproduction and increase its costs (English et al., 2012). Reproducing seasonally in an unpredictable environment, particularly when the latter is unproductive, could indeed lead to reproductive failures due to an unexpected period of food scarcity during the reproductive season, as well as to missed reproductive opportunities in case of an unexpected food burst outside the breeding season. In such cases, flexible reproductive schedules should be favoured. Additionally, following reproductive failures due to unexpected climatic events, non-seasonal breeders would be able to reproduce again rapidly, while strictly seasonal breeders would pay extra costs by having to wait for the next reproductive season.

Finally, and along the same line, a high rate of offspring mortality relative to adult mortality would amplify the costs of reproductive failures by multiplying their occurrence, without impacting the survival of adult females that would experience an extended gap before their next reproduction. Infanticide by males as well as by females, as a cause of mortality affecting young but not adults, might therefore contribute to favour the evolution of non-seasonal breeding. Infanticide by males is already known to be associated with non-seasonal breeding (Lukas and Huchard, 2014), the most common explanation positing that infanticide by males is unlikely to evolve in a context of seasonal breeding, because mothers of killed infants would not become sexually available to the killer male before the next breeding season, delaying the reproductive benefits of infanticide, so that non-seasonal reproduction would necessarily precede its appearance. Here we propose an alternative scenario positing that non-seasonal breeding and infanticide may instead have co-evolved, in a self-reinforcing loop.

### Objectives

In this study, we test this theoretical framework through a comparative analysis of a dataset with high-resolution quantitative measures of reproductive seasonality from a monophyletic taxonomic group, the order Primates. With their strong diversity of reproductive seasonality despite living in tropical environments (e.g. Campos et al., 2017), primates constitute a well-suited order to investigate the eco-evolutionary drivers of reproductive phenology. Contrary to higher latitudes, environmental productivity is high and environmental seasonality is less marked in the intertropical belt, which should alleviate the constraints leading to the evolution of seasonal reproduction, therefore providing an ideal environmental context to investigate other, overlooked determinants of reproductive seasonality. Additionally, primates have been extensively studied, providing rich datasets on wild populations (e.g. Trébouet et al., 2021 in the *Macaca* genus; Dezeure et al., 2024 in Africa-dwelling papionins), even though no large-scale comparative study has yet been conducted.

We tested the nine hypotheses presented above and listed in Table 1, first including the traditional hypothesis stating that reproductive seasonality is more frequent at high latitudes (H1). Second, our framework (see Fig. 1) predicts that reproductive decisions are limited by available resources. We therefore assessed the effect of environmental productivity, an index of baseline resource availability, where a greater productivity is expected to reduce reproductive seasonality (H2). Third, we expected environmental seasonality to increase reproductive seasonality (H3). Next, we designed an index of the intensity of daily reproductive effort, where a more intense reproductive effort is expected to favour the emergence of seasonal breeding (H4). We then investigated if three potential buffering mechanisms of environmental seasonality may reduce the intensity of reproductive seasonality, including the number of foraging innovations (Cognitive buffer – H5), diet breadth (Ecological buffer – H6) and allomaternal care (Social buffer – H7). Fourth, we expected environmental unpredictability to decrease reproductive seasonality (H8). In all these hypotheses based on the maintenance of positive energetic budgets (H1-8), we included environmental productivity as an additional predictor in order to control for cross-species variation in baseline resource availability. Finally, our last hypothesis states that seasonal breeding is counter-selected in cases where offspring mortality largely exceeds adult mortality, because females need to resume fertility rapidly after infant loss to minimize reproductive gaps (H9); infant mortality was here indexed by species exposure to infanticide.

**Table 1:**
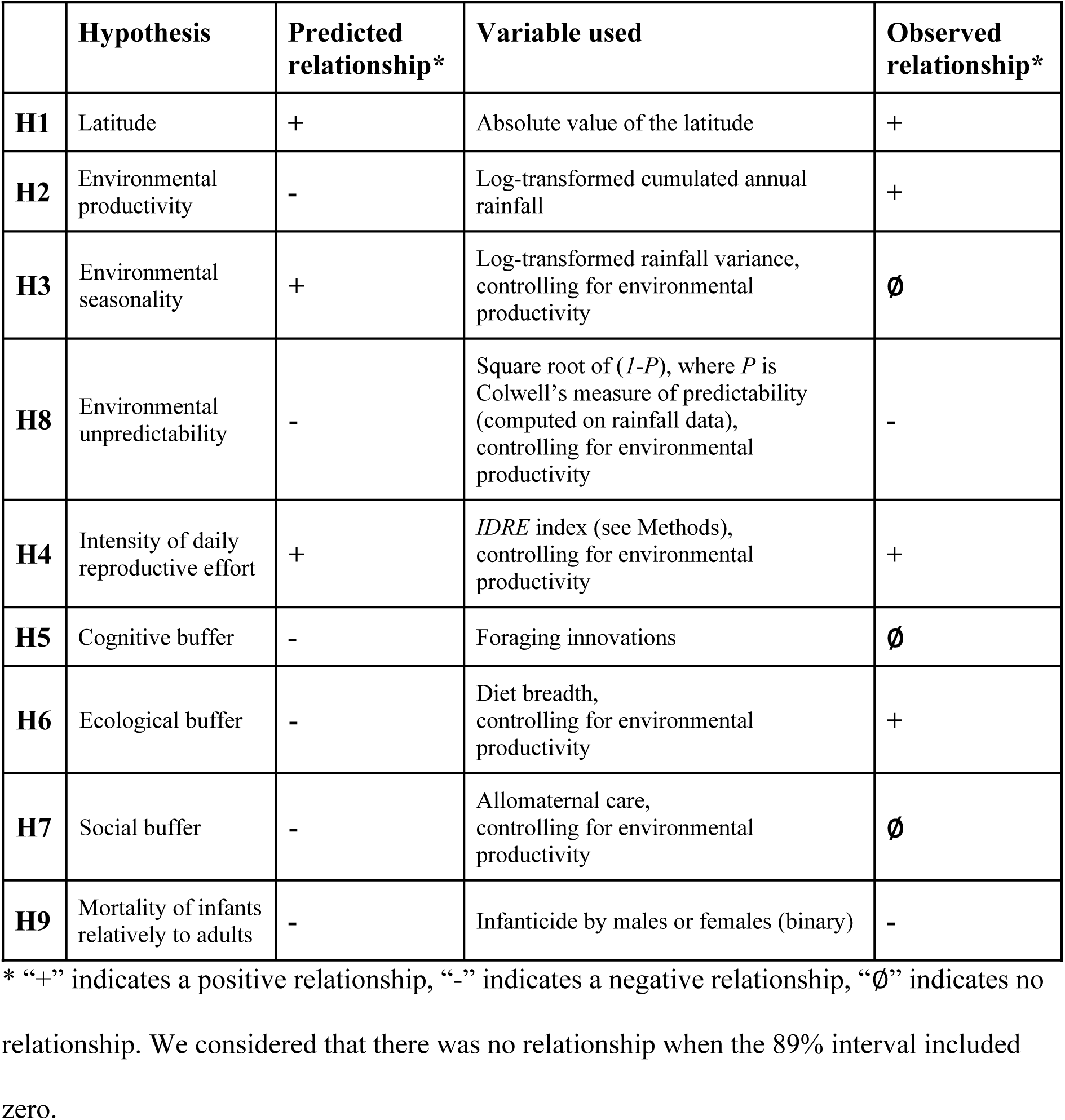
Summary of the hypotheses, predictions, and results of our tests.

## Methods

### Variables used

#### 1 Reproductive seasonality

A broad literature search was performed to gather high resolution data on reproductive seasonality in primates (Table S1). We looked for raw data on birth distributions in natural conditions for each primate species. We started with references listed in previous reviews (e.g. Heldstab et al., 2021). We then searched for species-specific data using Google Scholar with the following entries: “birth” OR “reproduction” and the name of the species, based on the list of primate species from the Mammal Diversity Database (Burgin et al., 2018). We chose Google Scholar as the database because data on reproductive timing is sometimes included in books or reports that are not covered in databases that focus on scientific literature. We used alternative or old species names in order to facilitate datasets matching for the analyses (Table S2). For each species, we recorded birth distributions for every population with available data, as the sample size did not affect the intensity of reproductive seasonality (*n* ranging between 4 and 7402, *rho* = 0.04, *p* = 0.68; Fig. S1). When several studies were available for the same population, we used data from the study with the highest number of births reported. We only collected data from non-provisioned, wild populations. When necessary, we extracted numerical data from graphs using WebPlotDigitizer (Drevon et al., 2017). We computed, for each monthly distribution of births, a quantitative measure of reproductive seasonality using circular statistics, according to the most recent recommendations (Thel et al., 2022) and considering that each birth occurred in the middle of the month (i.e. the 15^th^ of each month). In circular statistics, days of the year are represented as angles on a circle representing the annual cycle, and each birth event is characterised by a vector of length 1 pointing to the day of year when it occurred. By computing the mean vector for all birth events, it is possible to characterise their seasonality: the direction (*µ*) of the mean vector represents the mean date of birth, while its length (*r*) defines the intensity of seasonality (*r* = 0 when births are evenly distributed and *r* = 1 when all births occur on the same day). When the matching quantitative measures of reproductive seasonality (*r*) were given in the literature but the raw data on births was unavailable, we directly used the quantitative measure.

#### 2 Environmental variables

For each primate population included, the latitude of the study site was extracted during the literature search, together with data on reproductive seasonality. At this broad scale, commonly used (though imperfect) proxies for food availability rely on the amplitude of vegetation variation, which is primarily associated with climatic factors such as precipitation and temperature. Precipitation can accurately predict vegetation growth in tropical habitats (Nemani et al., 2003; Wu et al., 2015) and is therefore regularly used as a proxy of food availability when studying non-human primates (e.g. Dezeure et al., 2024). We used the coordinates of each study site to retrieve the monthly rainfall between January 1980 and December 2020 from the CHELSA database (https://www.chelsa-climate.org/, Karger et al., 2025), using the package *Rchelsa* (Karger, 2025). During preliminary checking, we removed *n* = 62 outlier values of monthly rainfall > 1250 mm (i.e. 0.03% of the initial dataset). Environmental productivity was measured as the log-transformed cumulative annual precipitation value averaged across 1980-2020; environmental seasonality was measured as the log-transformed average intra-annual variance: *environmental seasonality = log(mean(var(rainfall_m,y_)_y_))*, where *m* corresponds to the month and *y* corresponds to the year. Environmental unpredictability was measured as the square root of *(1-P)*, where *P* is Colwell’s measure of predictability (Colwell, 1974), an information-theory-based index that ranges between 0 (completely unpredictable) and 1 (fully predictable). This index of predictability is widely used in the literature (e.g. Loe et al., 2005; English et al., 2012; Kauffert et al., 2025) and has two components, constancy (*C*) and contingency (*M*), with *P = M + C*. Constancy reflects cases where a variable is predictable because its value remains constant throughout the year, while contingency captures cases where a variable follows the same, regular pattern of variation each year.

#### 3 Intensity of daily reproductive effort

The intensity of daily reproductive effort can be conceptualized by the amount of energy per time unit that a female needs to sustain reproduction (*E_repro_*; Hamilton et al., 2011). Its main part is used for the growth of offspring and therefore correlates with the overall offspring body mass that is created between conception and the end of lactation:

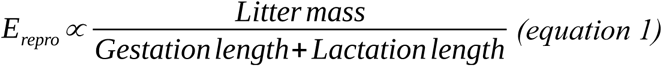

where *Litter mass* corresponds to the product of *Litter size* and *Weaning mass*. To assess the importance of the energetic cost associated with this amount of energy, it needs to be compared with a non-reproductive basal energy reference (*E_basal_*). Basal metabolism is known to be correlated to the 3/4 power of body weight (White et al., 2019) and can therefore be described as follows:

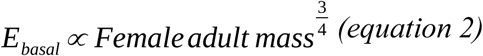

We therefore defined an index of the intensity of the daily reproductive effort (*IDRE*) that correlates with 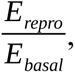 as follow:

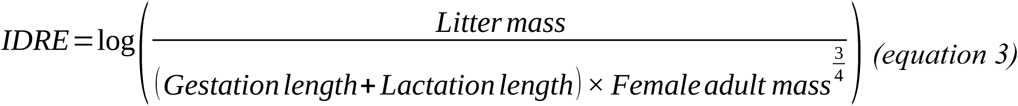

where values of *Litter size*, *Weaning mass*, *Gestation length*, *Lactation length* and *Female adult mass* were taken from Myhrvold et al. (2015) as well as Chiou et al. (2019), Schneider-Crease et al. (2022) and Weyher (2024) for the recently described kinda baboon (*Papio kindae*). Additional values of female adult mass were taken from West and Capellini (2016), using the mean value when both studies gave different values for the same species. Because this index is linearly distributed, we log-transformed it to obtain an allometric scaling, more suited to variables involving body mass (Anderson-Teixeira et al., 2009).

#### 4 Foraging innovation (cognitive buffer)

The number of innovations that occur in a foraging context was extracted from Navarrete et al. (2016) based on a previous study computing examples of innovation, social learning, tool use and extractive foraging in living non-human primates (Reader et al., 2011). The number of published articles by species was also extracted and used to control for search effort.

#### 5 Diet breadth (ecological buffer)

Data on diet was extracted from Machado et al. (2022), this study itself based on the diet information provided by Mittermeier et al. (2013). The authors designed a diet richness index representing the number of categories eaten by a species from a list of 40 different food items (Machado et al., 2022).

#### 6 Allomaternal care (social buffer)

Data on allomaternal care was extracted from Heldstab et al. (2019), which complements the databases of Isler and van Schaik (2012) and Heldstab et al. (2017). In this study, allomaternal care (by males and by other group members) is described as a continuous variable based on the frequency of occurrence of provisioning, carrying, protection, and other energetically influential care behaviours (e.g. huddling, communal nesting and pup retrieval), excluding allonursing (Heldstab et al., 2019).

#### 7 Infanticide

Data on infanticide by males and by females (binary variable reported as “yes” or “no”) were extracted from Lukas and Huchard (2014) and Lukas and Huchard (2019) (and from Schneider-Crease et al. (2022) for kinda baboons) and later combined into a single variable describing whether or not infanticide of any sort (by males and/or by females) was observed.

### Phylogeny

We downloaded a credible set of 1000 trees of mammalian phylogenetic history from vertlife.org/phylosubsets/ (June 2023) and used TreeAnnotator (version 1.10.4 in BEAST; Drummond et al., 2012) to generate a maximum clade credibility (MCC) tree (median node heights and burn in of 250 trees). We trimmed the tree to match the species in our sample (on a limited number of occasions, where our sample species was not represented in the source phylogeny, we used a sister taxon). We tested the strength of the phylogenetic signal in reproductive seasonality among each dataset using Blomberg’s *K*, where *K* values close to 0 means that closely related species are not similar to each other and *K* values close to 1 means that they are. This test was associated with a p-value (based on 1000 randomizations) assessing whether closely related species are more similar to each other than expected by chance.

### Analyses

We performed all analyses in the statistical software R (version 4.5.2; R Core Team, 2025). For each predictor variable, we excluded from the respective analysis species for which we could not find data (we report the sample size for each analysis). We used a Bayesian approach to build statistical models and estimated relationships as implemented in the package *rethinking* using the function *ulam* (McElreath, 2020) to fit the models with Markov chain Monte Carlo estimation in stan (Stan Development Team, 2025). We fitted multilevel models that include the shared phylogenetic history as a covariance matrix. Weakly regularizing priors were used for all parameters, and we drew 1000 samples from four chains. These posterior samples from each model were used to generate estimates of the overall effect of each predictor. We determined whether a variable had a relationship with the variation in reproductive seasonality when the 89% compatibility estimate of the posterior sample did not cross zero (continuous predictor variable: latitude, productivity, seasonality, unpredictability, intensity of daily reproductive effort, diet breadth and allomaternal care; all standardized by subtracting the mean and dividing by the standard deviation) or if the contrast between levels did not cross zero (categorical predictor variable: infanticide), indicating that our data showed a consistent positive or negative association. The response variable of the phylogenetic multilevel models was the reproductive seasonality of the species or populations considered, a standardised continuous variable, except for the model investigating the relationship between reproductive seasonality and number of foraging innovations. In this case, we used the number of foraging innovations as the response variable and the seasonality of births as the explanatory variable, along with the number of published articles by species as an additive effect. This approach was chosen because the number of foraging innovations reported for a given species is biased by search effort, as the more a species has been studied, the higher the chances to document a new foraging behaviour (Navarrete et al., 2016).

We first checked which of the environmental variables (productivity, seasonality and unpredictability) was most directly associated with the variation in reproductive seasonality using univariate models and found that environmental productivity had the strongest explanatory power. We therefore used bivariate models including environmental productivity as a control variable when testing the relationship between reproductive seasonality and the predictors that are expected to show conditional effects (environmental seasonality, environmental unpredictability, daily reproductive effort, diet breadth and allomaternal care). Considering the different approach taken in this model, we could not add environmental productivity as a control variable when testing the number of cognitive innovations. We used univariate models when testing the relationship between reproductive seasonality and the predictors that are expected to show direct effects (environmental productivity, latitude and infanticide).

We estimated *ɑ* as the analytical mean (or intercept) of the reproductive seasonality values, *β*, the estimate of the predictor, for which we determined if its compatibility interval crossed zero, and, when appropriate, *γ*, the estimate of the control variable (environmental productivity). The model in *rethinking* took the form:

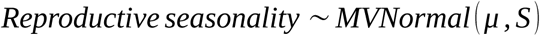

Where

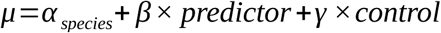

With

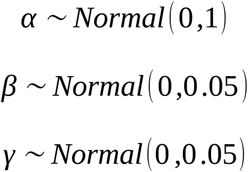

And

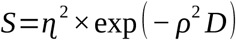

with

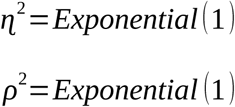

where the *Reproductive seasonality* values come from an overall distribution of mean *μ* and of variance-covariance matrix *S*. *μ* depends on the mean *ɑ* and the relationship with the respective additive predictor variables *β × predictor* and *γ × control*. The priors for *β* and *γ* are centered around zero assuming that the standardized predictor and control variable might have no, a negative, or a positive influence. *S_ki,lj_* reflects the similarity between observation *i* of the species *k* and observation *j* of the species *l*. *S* is the variance-covariance matrix of *Reproductive seasonality*, where the same species *k* can appear in multiple rows or columns when there are multiple populations with observed reproductive seasonality from that species. *S* is a transformation of the squared phylogenetic distance *D* among all species pairs *k*, *l*, which is the sum of the branch lengths separating species pairs *k* and *l* in the phylogeny, according to a Gaussian process with a multinormal prior with the parameters *ɳ*^2^ (maximum covariance among closely related species) and *ρ*^2^ (decline in covariance as phylogenetic distance increases), whose priors are constrained to be positive. The only difference for the categorical predictor variables is that, instead of estimating a linear relationship *β × predictor*, we estimate the separate intercepts for the different categories *β_c_* [*predictor*].

## Results

### Data description

We collected *r* values of reproductive seasonality for 132 populations from 94 primate species during our literature search (Fig. 2, Table S2). Their spatial repartition ranged between latitudes 34°27’S and 36°27’N (Fig. S4, Fig. S5). *Cercopithecidae* represented 52% of the populations sampled, while they represent only 33% of all primate species. Conversely, *Cheirogaleidae* (42 species), *Lepilemuridae* (25 species) and *Pitheciidae* (56 species) were underrepresented with respectively zero, zero and two populations in our dataset (Fig. S6). Nevertheless, *Lemuridae* (21 species), as well as *Atelidae* (23 species) and *Cebidae* (30 species) were well represented with respectively 11, 13 and 11 populations. There was a significant, but small phylogenetic signal for reproductive seasonality among the dataset (Blomberg’s *K* = 0.12, *p* < 0.01). This reflects that, while some clades are relatively consistent (e.g. all lemurs are relatively seasonal), the extent of seasonality could differ dramatically among closely related species in other clades (e.g. *Macaca* genus, Fig. 2).

**Figure 2:**
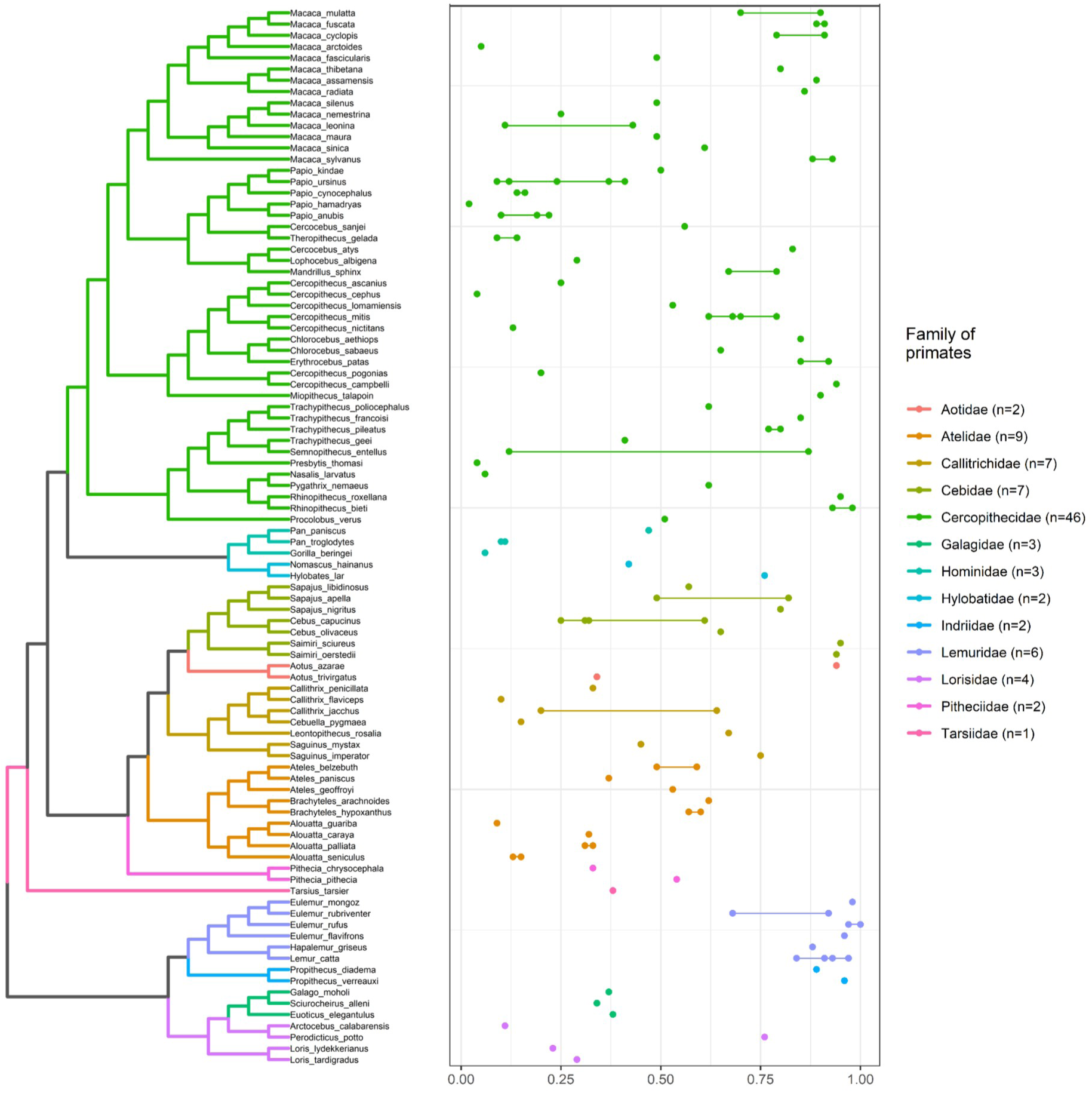
Phylogenetic distribution of the *r* values of reproductive seasonality ranging from 0.02 for the least seasonal species, hamadryas baboon (*Papio hamadryas*) to 1.00 for the highest seasonality, red-fronted lemur (*Eulemur rufus*). In this graph, *r* values of reproductive seasonality were averaged when several populations of the same species were available. About half of the species sampled were *Cercopithecidae* (46 out of 94 species). Reproductive seasonality varied widely among almost all primate families with the exception of lemurs (*Lemuridae* and *Indriidae*) that were highly seasonal.

### Test of hypotheses

#### 1 Support for the ecological hypotheses (H1-3, 8)

Latitude (absolute distance to equator) was positively associated with the intensity of reproductive seasonality (Fig. 3 and Fig. 4A, 89% interval: [0.01; 0.12]). When investigated with univariate models, the three environmental variables were positively associated with reproductive seasonality (environmental productivity 89% interval: [0.17; 0.23]; environmental seasonality 89% interval: [0.16; 0.22]; environmental unpredictability 89% interval: [0.13; 0.19]). When both environmental productivity and seasonality were included in the same bivariate model, environmental productivity (89% interval: [0.11; 0.22]) had a stronger coefficient than environmental seasonality (89% interval: [-0.02; 0.10]). We found similar results with environmental productivity (89% interval: [0.37; 0.51]) and environmental unpredictability (89% interval: [-0.35; -0.20]).

**Figure 3:**
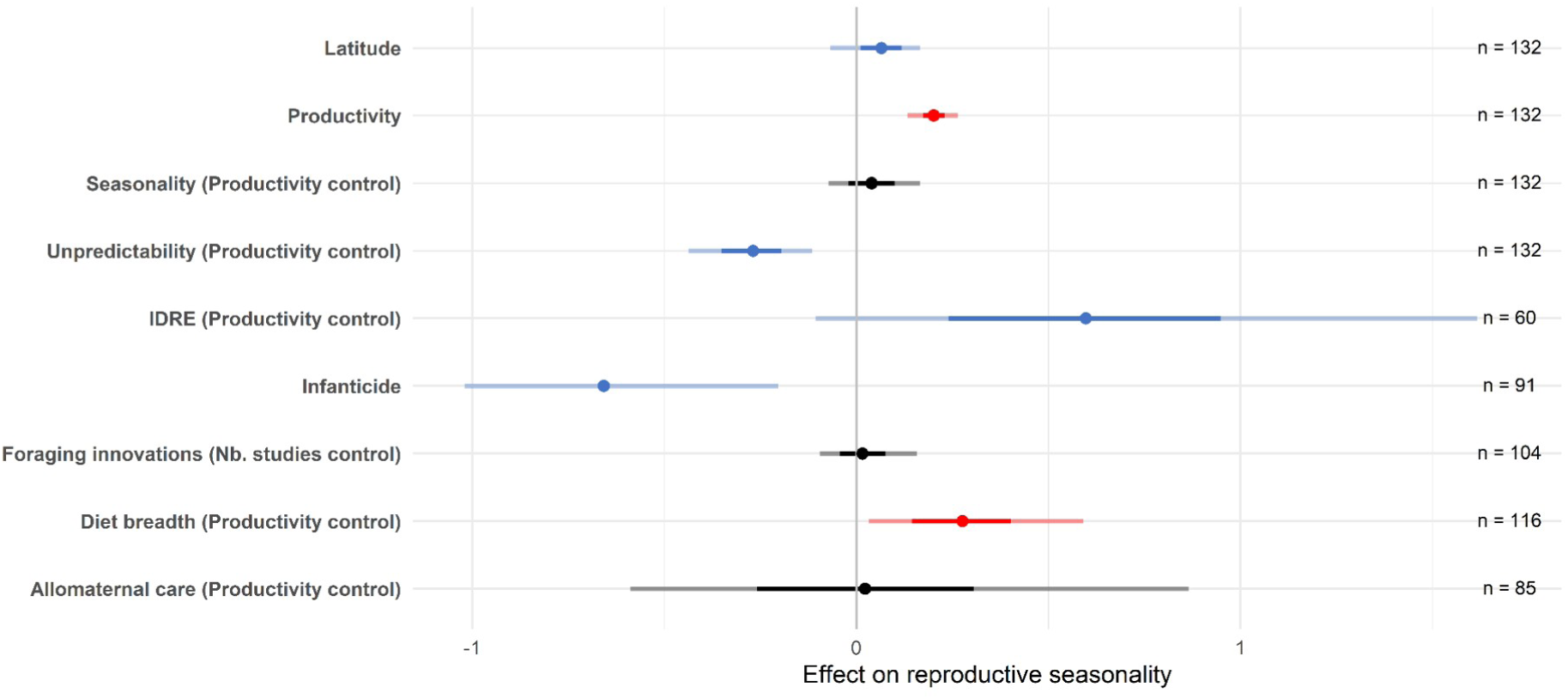
Main predictors of reproductive seasonality. Dots represent the estimated mean effects, bold lines represent the 89% interval of the mean effects, thinner lines represent the range of the prediction estimates of the models. Blue and red colours indicate a significant relationship (89% interval do not include zero), either in accordance with our predictions (blue) or in disagreement (red). Black indicates a non-significant relationship (89% interval includes zero). Models are univariate unless specified otherwise (bivariate models with a control variable for environmental productivity or the number of studies). The number of populations for which data were available are provided. IDRE stands for the intensity of daily reproductive effort.

**Figure 4:**
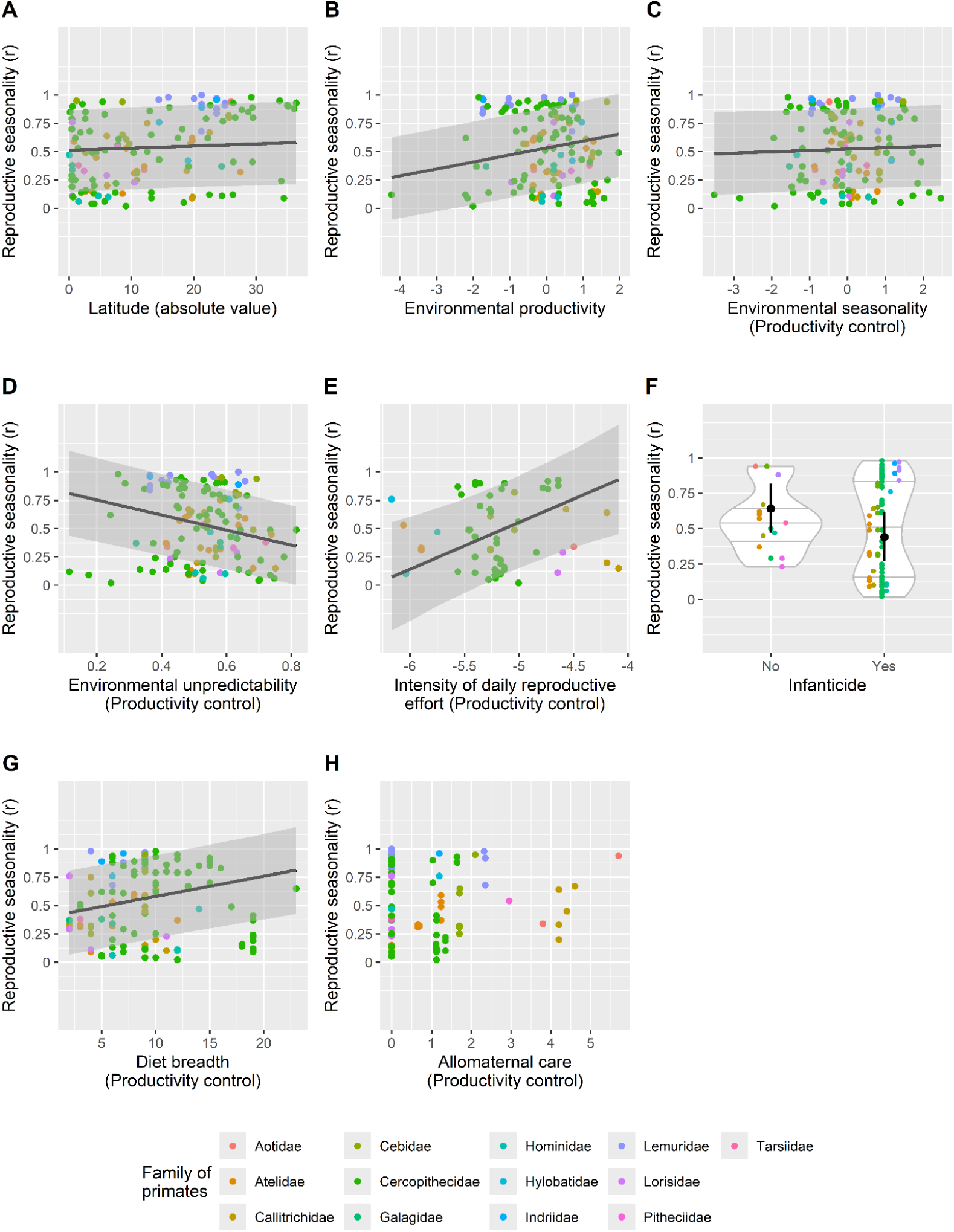
Relationship between the main predictors investigated (A-H) and reproductive seasonality. Dots represent the raw values observed for each population, their colours indicating the associated family of primates. For a better visualisation, points are dodged when overlapping. For predictors with a significant association with reproductive seasonality, the black lines represent the relationship as predicted by the *rethinking* models (for an averaged environmental productivity as a control variable, when appropriate, see text for detail), with grey areas (black segments, when appropriate) showing the 89% interval.

However, unexpectedly, environmental productivity (measured as the log-transformed mean cumulative annual rainfall) was positively associated with seasonality of reproduction (Fig. 3 and Fig. 4B, 89% interval: [0.17; 0.23]). When controlling for environmental productivity, environmental seasonality (measured as the log-transformed variance of monthly rainfall across the annual cycle) was marginally positively associated with reproductive seasonality (Fig. 3 and Fig. 4C, 89% interval: [-0.02; 0.10]), and environmental unpredictability (measured as the square root of (*1-P*) where *P* is Colwell’s measure of predictability) was negatively associated with reproductive seasonality (Fig. 3 and Fig. 4D, 89% interval: [-0.35; -0.20]).

#### 2 Support for the daily reproductive energy expenditure hypothesis (H4)

A strong positive relationship was observed between the intensity of daily reproductive effort and reproductive seasonality, both when controlling for environmental productivity (Fig. 3 and Fig. 4E, 89% interval: [0.24; 0.95]) or not (89% interval: [0.09; 0.76]).

#### 3 No support for the buffer hypotheses (H5-7)

We did not identify a significant relationship between allomaternal care and reproductive seasonality, either when controlling for environmental productivity (Fig. 3 and Fig. 4H, 89% interval: [-0.26; 0.30]) or not (89% interval: [-0.25; 0.30]). Similarly, we did not identify a significant relationship between the number of foraging innovations observed in a primate species and reproductive seasonality (Fig. 3, 89% interval: [-0.04; 0.07]). The extent of diet breadth was widely variable, with four species (black tufted-ear marmoset (*Callithrix penicillata*), moholi bushbaby (*Galago moholi*), slender loris (*Loris tardigradus*) and Bosman’s potto (*Perodicticus potto*)) displaying the smallest diet breadth with a score of 2, and one species (green monkey (*Chlorocebus sabaeus*)) displaying the largest diet breadth with a score of 23. Contrary to our expectations, diet breadth was positively associated with reproductive seasonality, both when controlling for environmental productivity (Fig. 3 and Fig. 4G, 89% interval: [0.14; 0.40]) or not (89% interval: [0.14; 0.38]).

#### 4 Support for the mortality hypothesis (H9)

We detected a strong, significant negative relationship between the occurrence of infanticide from males and/or females and reproductive seasonality (Fig. 3 and Fig. 4F, 89% interval: [-0.84; -0.47]). We found very similar results when considering infanticide committed only by males (89% interval: [-0.85; -0.47]) and only by females (89% interval: [-0.80; -0.45]).

## Discussion

### Summary of the results

We developed a theoretical framework for explaining seasonal breeding in mammals and tested it on 132 populations of 94 species of primates. As expected, we found that reproductive seasonality increases when latitude (H1), environmental seasonality (H3) and environmental predictability (H8), when controlling for environmental productivity.

However, contrary to our expectations, reproductive seasonality increases with environmental productivity (H2), with which it is more strongly associated than the other environmental variables. Such results suggest that at the relatively low latitudes inhabited by most primates, productivity plays a stronger role than seasonality in shaping life history, potentially indicating that among primates, reproductive seasonality does not occur as an adaptive response to limit reproduction during seasonal droughts, but as an adaptive response to accelerate reproductive pace during times of abundance. As expected, reproductive seasonality increases with daily reproductive effort (H4), indicating that challenges associated with financing reproductive efforts are a major driver of life history pace. Contrary to our expectations, there is no evidence of the role of buffering strategies (cognitive buffer, H5; social buffer, H7; ecological buffer, H6) in releasing the necessity for a seasonal reproduction in primates. Our framework assumed that buffer strategies might be needed when species face limitations in resource availability, but in our results, the unexpected effect of environmental productivity suggests that such limitations might not be the key factor shaping reproductive timing among primates living almost exclusively within the tropics. Lastly, reproductive seasonality is negatively correlated with the occurrence of infanticide by males or by females (H9), suggesting that non-seasonal breeding and infanticide may have co-evolved in a self-reinforcing loop. These findings provide new insights for our understanding of reproductive seasonality in mammals, including humans.

### Reproductive seasonality is common in productive environments (H1-3, 8)

We detected a positive effect of latitude on reproductive seasonality (H1), confirming many previous studies in mammals (in ruminants, Zerbe et al., 2012; in Carnivora, Heldstab et al., 2018; in lagomorphs, Heldstab, 2021a; in rodents, Heldstab, 2021b; in primates, Heldstab et al., 2021). Despite living in relatively constrained latitudes—mostly within the tropics—primates inhabiting higher latitudes are significantly more seasonal.

Environmental productivity decreases with increasing distance to the equator (Fig. S7), but contrary to our expectations and a previous study (Burtschell et al., 2023), we found a positive effect of environmental productivity on reproductive seasonality (H2). This result may reflect a genuine positive relationship between environmental productivity and reproductive seasonality. In line with this, a previous study on Africa-dwelling papionins also found such a positive relationship and suggested that reproductive failures might occur more frequently in low-productivity environments, thus contributing to select for non-seasonal breeding in order to enhance female’s ability to resume cycling rapidly following such failures (Dezeure et al., 2024). In more productive environments, selection may promote higher reproductive investment per unit time, leading to faster weaning and a reduced probability of reproductive failure. In line with this, mandrills (*Mandrillus sphinx*) breed seasonally while living in the equatorial forest, and display an unusually fast reproductive pace, where females can conceive again while they are still lactating (Dezeure et al., 2023). In summary, building on Dezeure et al. (2023, 2024) and the present study, hypothesis H2 likely requires reformulation. In species with relatively slow reproduction, reduced environmental productivity may select against reproductive seasonality by favouring strategies that minimize the reproductive gaps following reproductive failure, whereas increased productivity may select for faster reproductive pace, and in turn reproductive seasonality. Under such conditions, reproductive seasonality may reduce vulnerability to reproductive failure by preventing failure from occurring, rather than by compensating for them once they have occurred.

Although surprisingly marginal, the effect of environmental seasonality, of which latitude is often considered to be the proxy (even though this is not the case in our dataset; Fig. S7), is visible in our results but non-significant after controlling for environmental productivity (H3), suggesting that its effect on reproductive seasonality is not as strong as previously thought in primates. Nevertheless, even though tropical environments are often considered to display moderate seasonal variation at macro-ecological scales, food resources in these systems can, in fact, be markedly seasonal (Sakai, 2001; Feng et al., 2013). Since most primates selectively feed on certain plant species, their reproductive phenology can be affected by the seasonal cycle of the species that form the basis of their diet (Di Bitetti and Janson, 2000). For example, kinda baboons, who are omnivorous with a dominance of frugivory, are the only baboon species that breeds seasonally (Petersdorf et al., 2019). Living further away from the equator might also expose some species to additional seasonal climatic fluctuations more directly, something our analyses did not focus on. For example, individuals might time their reproduction in response to extreme temperatures if young offspring are sensitive to low or high temperatures (Zhao et al., 2020; Khera et al., 2023).

Our study provides support for the expected negative effect of environmental unpredictability on reproductive seasonality (H8), as we found a negative relationship between these two variables after controlling for environmental productivity. Environmental unpredictability is positively correlated with environmental productivity—and a cortege of environmental variables—in our dataset (Fig. S2), so that confounding effects likely mask the negative relationship when these correlations are not accounted for. Unpredictable but highly productive systems are likely less constraining than unpredictable and low productivity systems, where it becomes less advantageous to reproduce seasonally. However, a number of unpredictable systems are also relatively productive in the primates’ distribution range (Fig. S2), which leaves only a limited number of “harsh” systems to test the full range of environmental conditions experienced by mammals. The mixed effects regarding the association between environmental unpredictability and breeding seasonality detected in previous studies (Loe et al., 2005; English et al., 2012; Dezeure et al., 2023, 2024; Kauffert et al., 2025) could be due to the fact that the metrics used to describe environmental unpredictability vary between studies, but also and above all, that these approaches do not control for other environmental parameters. Future research efforts should design quantitative indices of predictability that are less dependent on other environmental variables. More generally, the correlations between different explanatory variables should be accounted for with multivariate models when there is enough variability in the data.

Overall, our results underline the complexity of environmental pressures shaping reproductive seasonality in tropical mammals. A classic example of tropical mammals displaying intense reproductive seasonality is the monophyletic group of lemurs, where females display intense reproductive seasonality (Wright, 1999; Kappeler and Fichtel, 2015), the determinants of which are not fully elucidated (Heldstab et al., 2021). Here we found no clear evidence that the climatic variables characterizing Malagasy study sites are more seasonal, more productive or less unpredictable than for the rest of the dataset (Fig. S8); in fact, lemurs tend to occupy intermediate positions on those variables. These findings fail to support a primary role of the Malagasy climate and may instead point to the role of their particular life-history (with a relatively fast reproductive pace compared to non-strepsirrhines; Harvey and Clutton-Brock, 1985) and physiological particularities (with their low resting metabolic rate; Ross, 1992) in affecting their reproductive strategies.

### Spreading reproductive effort over extended periods allows primates to reproduce less seasonally (H4)

The positive relationship between the intensity of daily reproductive effort and reproductive seasonality (H4) is significant, with one of the largest effect sizes. This result is consistent with a modelling study simulating energy budgets in a long-lived primate, that showed that a slight increase in offspring’s daily growth rate drastically increases the selective pressure for seasonal reproduction (Burtschell et al., 2023). The effect of several variables linked with our index (litter size, number of litters per year, gestation length, weaning age, adult body mass) has been investigated separately in previous studies (Zerbe et al., 2012; Owen-Smith and Ogutu, 2013; Heldstab et al., 2018; Heldstab, 2021a, 2021b; Heldstab et al., 2021).

Individually, most of them had no effect on the intensity of birth seasonality (with a few exceptions for some taxonomic groups that were all consistent with the positive relationship identified between our index of daily reproductive effort intensity and reproductive seasonality in primates). The fact that we detected a robust relationship with our index indicates that it’s the daily reproductive effort, and ultimately balancing daily energetic budgets, that is important in shaping female phenology strategies, more than the length of the different stages of the reproductive cycle.

More generally, this result emphasizes the crucial importance of life history variation in shaping patterns of reproductive seasonality, above and beyond environmental variables. It also explains why many long-lived species breed non-seasonally: for example, in great apes, infants are weaned at 5-6 years of age so that lactation covers five dry seasons, meaning that reproductive seasonality is not adaptive in this context. It also provides powerful support to the idea that adaptation to fluctuating resource availability is a key driver of life-history pace through reduced reproductive effort (Stearns, 1976; Jones, 2011), where females need to spread their reproductive costs over extended periods—thus slowing down offspring development and life-history—in order to cope with environmental variability.

### No evidence that behavioural strategies buffering the seasonal food shortage allow primates to reproduce less seasonally (H5-7)

We next tested whether primates mitigate seasonal food shortages through three behavioural strategies that could allow them to extract more energy during the lean season: innovating during foraging (cognitive buffer, H5), broadening the diet (ecological buffer, H6), or sharing reproductive costs across group members (social buffer, H7). We found no support for these hypotheses. Several interpretations are possible. In light of our other findings on the environmental and life-history drivers of breeding seasonality, one possibility is that reproductive seasonality in primates evolves more in response to environmental opportunities than to environmental constraints. Most primates have interbirth intervals that are too long to fit within an annual cycle, preventing them from buffering energetic bottlenecks through regular “annual pauses” in reproduction. Flexible reproduction may instead allow them to capitalize on seasonal food peaks to decrease interbirth intervals, minimize offspring mortality, or offset the costs of infant mortality by rapidly resuming fertility (Dezeure et al., 2021, 2023, 2024). In this context, while behavioural buffers may be essential to survive and reproduce during the lean season, they do not shape the evolution of their reproductive phenology, as we initially predicted.

The positive association between dietary breadth and breeding seasonality supports this “opportunities/plasticity” interpretation. Generalist species, who are well equipped to be ecologically opportunistic, may be especially able to exploit seasonal food peaks. Comparing interbirth intervals (IBIs) preceding synchronized versus non-synchronized births could help distinguish between the harshness/buffering and opportunities/plasticity models. The harshness/buffering model predicts longer IBIs before synchronized births, because females must delay reproduction until conditions improve. In contrast, the opportunities/plasticity model predicts shorter IBIs, as females accelerate reproduction in response to favourable conditions. Supporting the latter interpretation, a study in mandrills, where females can conceive year-round yet still display pronounced birth peaks, shows that only high-ranking female mandrills manage to fit their IBIs within a year and breed every year (Dezeure et al., 2024).

A second possibility is that our proxies of behavioural buffers are noisy. For example, in our study, allomaternal care includes not only energetically beneficial behaviours, but also behaviours such as protection, huddling and communal nesting (Heldstab et al., 2019). Additionally, it is cooperative breeding, more than allomaternal care, that has previously been found as a strategy to buffer environmental variations and cope with harsh or unpredictable environments in African birds (Rubenstein and Lovette, 2007) and mammals (Lukas and Clutton-Brock, 2017). Yet, cooperative breeding is associated with the production of litters rather than singletons, a trait that is rare in primates. Furthermore, the positive relationship between the number of studies and the number of foraging innovations indicates that our index of cognitive buffer might be underestimated among less studied species thus probably noisy (Navarrete et al., 2016).

### Infanticide promotes non-seasonal reproduction (H9), especially in slow-living species where adult mortality is low

In line with previous studies (Brockman and van Schaik, 2005; Lukas and Huchard, 2014), we found a strong negative relationship between infanticide (by males and/or by females) and reproductive seasonality (H9). When species reproduce slowly, a secondary increase in offspring mortality (relative to adult mortality) likely increases the costs of reproductive seasonality because non-seasonal breeders could initiate a new reproductive attempt immediately after the failed one, while seasonal breeders may have to wait for the next reproductive season. Indeed, the theory of sexually selected infanticide (Hrdy, 1979) posits that infanticide is only beneficial for males when it accelerates female’s return to fertility. Such benefits may be offset if females reproduce seasonally and have to wait for the next breeding season. While the literature has traditionally considered a unilateral relationship between infanticide and seasonal breeding (Brockman and van Schaik, 2005; Lukas and Huchard, 2014), positing that reproductive (non-)seasonality is a causal factor determining the evolution of infanticide by males, our approach, which considers infanticide from both males and females, opens the alternative possibility of a bilateral, more dynamical, relationship between both traits, where infanticide and reproductive seasonality would evolve jointly in a feed-back loop, with offspring mortality favouring the evolution of non-seasonal breeding, which would in turn promote infanticide by males. Although our study does not allow the establishment of causal relationships, it highlights the plausibility of a new evolutionary scenario that refines our understanding of the evolution of infanticide—which was considered as well established. Additionally, it points to a new, so far overlooked, potential determinant of mammalian reproductive phenology.

### Significance for the understanding of human reproductive phenology

Our findings support a shift in perspective: in slow-living primates, reproductive seasonality is not a default, necessary response to seasonal food scarcity, but rather a flexible, facultative, and opportunistic adaptation to environmental fluctuations. This model likely extends to humans. The fact that all great apes are capable of year-round breeding (Campos et al., 2017) suggests that non-seasonal reproduction probably co-evolved with very slow life histories in equatorial primates—as a consequence of reduced reproductive effort (Jones, 2011)—long before hominization. Consequently, non-seasonal breeding is unlikely to be a human-specific trait resulting from our exceptional ability to buffer environmental variation through foraging innovations, extractive technologies, agriculture or niche construction, as has been proposed in the anthropological literature (O’Brien and Laland, 2012; Emery Thomson, 2013). In line with this, we found no correlation between behavioural buffers and reproductive non-seasonality across primates, and generally no support for this buffer/harshness model.

Furthermore, the reduction in food availability variance associated with the evolution of *Homo*—and the resulting more consistent access to food resources (Pontzer, 2012)—likely facilitated their colonization of diverse habitats, including low-productivity environments. Some degree of seasonal reproduction has indeed been documented across a range of human populations and environments (e.g., Turkana people, Kenya [Leslie and Fry, 1989]; pre-modern Finland [Lummaa et al., 1998]; modern Mexico [MacFarlan et al., 2021]; Baka Pygmies, Cameroon [Piqué-Fandiño et al., 2022]). In light of these results, humans—like other long-lived primates—may flexibly adjust their reproductive phenology to capitalize on food peaks, aligning with the opportunities/plasticity model we propose. Comparative studies of traditional human populations could further test this model, particularly the hypothesis that birth peaks are more pronounced in productive environments and among populations with more generalist subsistence strategies (controlling for latitude).

## Conclusion

This study sheds new light on the evolution of (non-)seasonal reproduction by proposing and testing an integrative theoretical framework. Nine hypotheses were tested by conducting comparative analyses in primates. Supporting previous work in this area, we confirmed that ecological factors, including environmental seasonality, productivity and unpredictability, play a major role in the evolution of seasonal breeding, though in unexpected ways. Our results indicate that in primates, reproductive seasonality has evolved in productive and predictable environments, as a flexible and facultative response to environmental opportunities, more than as a constraint to mitigate seasonal food scarcity. Another novelty of our work reveals the importance of life-history in shaping primate reproductive phenology. Specifically, the temporal spreading of reproductive effort, alongside substantial sources of infant mortality, are both associated with the evolution of non-seasonal breeding. Finally, our results fail to support hypotheses proposing that non-seasonal breeding reflects a behavioural capacity to buffer environmental variability, through cognition (foraging innovations), sociality (allomaternal care) or ecology (increased diet breadth). Altogether, these results support a shift in perspective in our understanding of the evolution of reproductive seasonality in primates that likely extends to humans as well as many slow-living tropical mammals. Instead of a default, necessary response to seasonal food scarcity, reproductive seasonality appears as a flexible, facultative, and opportunistic adaptation to environmental fluctuations.

## Authors’ contributions

Lugdiwine Burtschell: Conceptualization - Methodology - Software - Validation - Formal analysis - Investigation - Resources - Data Curation - Writing: Original Draft - Visualisation - - Funding acquisition.

Lucie Thel: Conceptualization - Methodology - Software - Validation - Formal analysis - Investigation - Resources - Data Curation - Writing: Review & Editing - Visualisation - Funding acquisition.

Dieter Lukas: Conceptualization - Methodology - Validation - Formal analysis - Investigation - Writing: Review & Editing - Visualisation - Supervision.

Jules Dezeure: Conceptualization - Methodology - Writing: Review & Editing. Bernard Godelle: Conceptualization - Methodology - Writing: Review & Editing.

Elise Huchard: Conceptualization - Methodology - Investigation - Resources - Writing: Review & Editing - Supervision - Project administration - Funding acquisition.

## Acknowledgments

We are very grateful to Marie Riffis, Suzie Tallon, Julien Offresson, Fantine Benoit and Amandine Merenciano for their help in collecting the data. Lugdiwine Burtschell thanks AgroParisTech and Lucie Thel thanks Nelson Mandela University for the financial support. This work was supported by the ANR grant “ERS-17-CE02-0008”, 2018-2021 awarded to Elise Huchard. Contribution ISEM n°[to_be_completed_upon_publication].

## Data availability statement

Data and scripts for analyses available from the Zenodo Repository: https://doi.org/10.5281/zenodo.20210295 (Anonymised, 2026).

