## supplementary material for "The evolution of non-seasonal breeding in primates"

**Figure S1**: Variation of reproductive seasonality according to the sample size of the populations of primates ($n$ = 132) included in the analyses. The population of rhesus macaque (*Macaca mulatta*) in Cayo Santiago is not visible in the figure ($n$ = 7402 births, $r$ = 0.70). A Spearman's rank correlation test showed no significant correlation ($rho$ = 0.04, $p$ = 0.68).

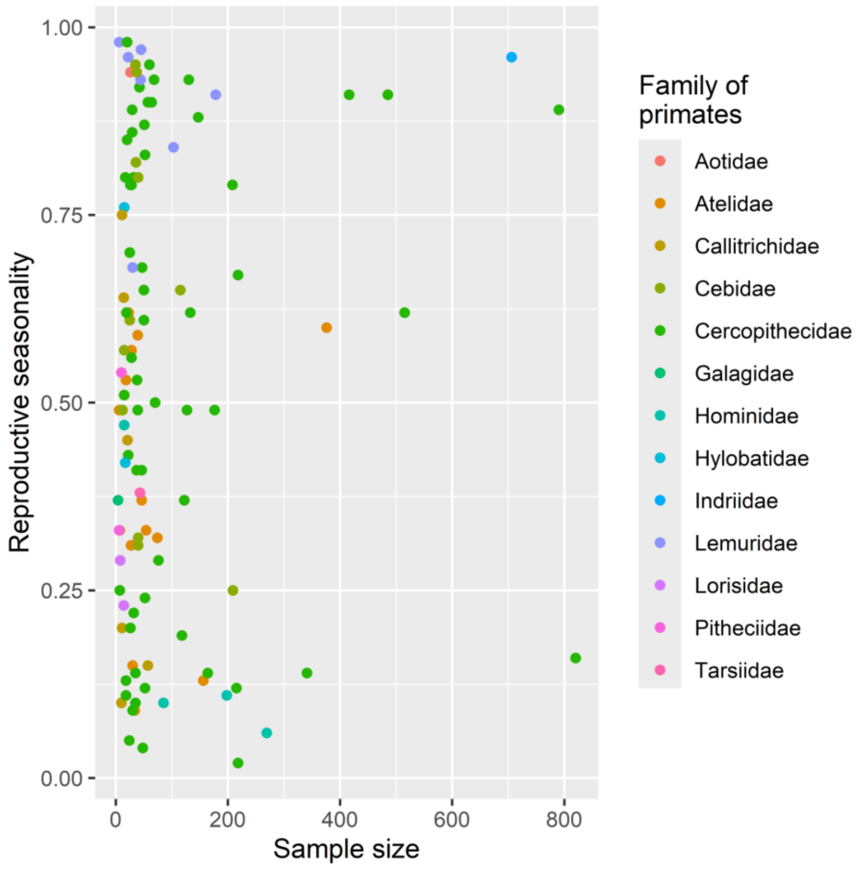

**Figure S2**: Covariation (Spearman's rank correlation *rho* and significance value *p*) among environmental productivity, seasonality and unpredictability (estimated based on monthly rainfall data between 1980 and 2020).

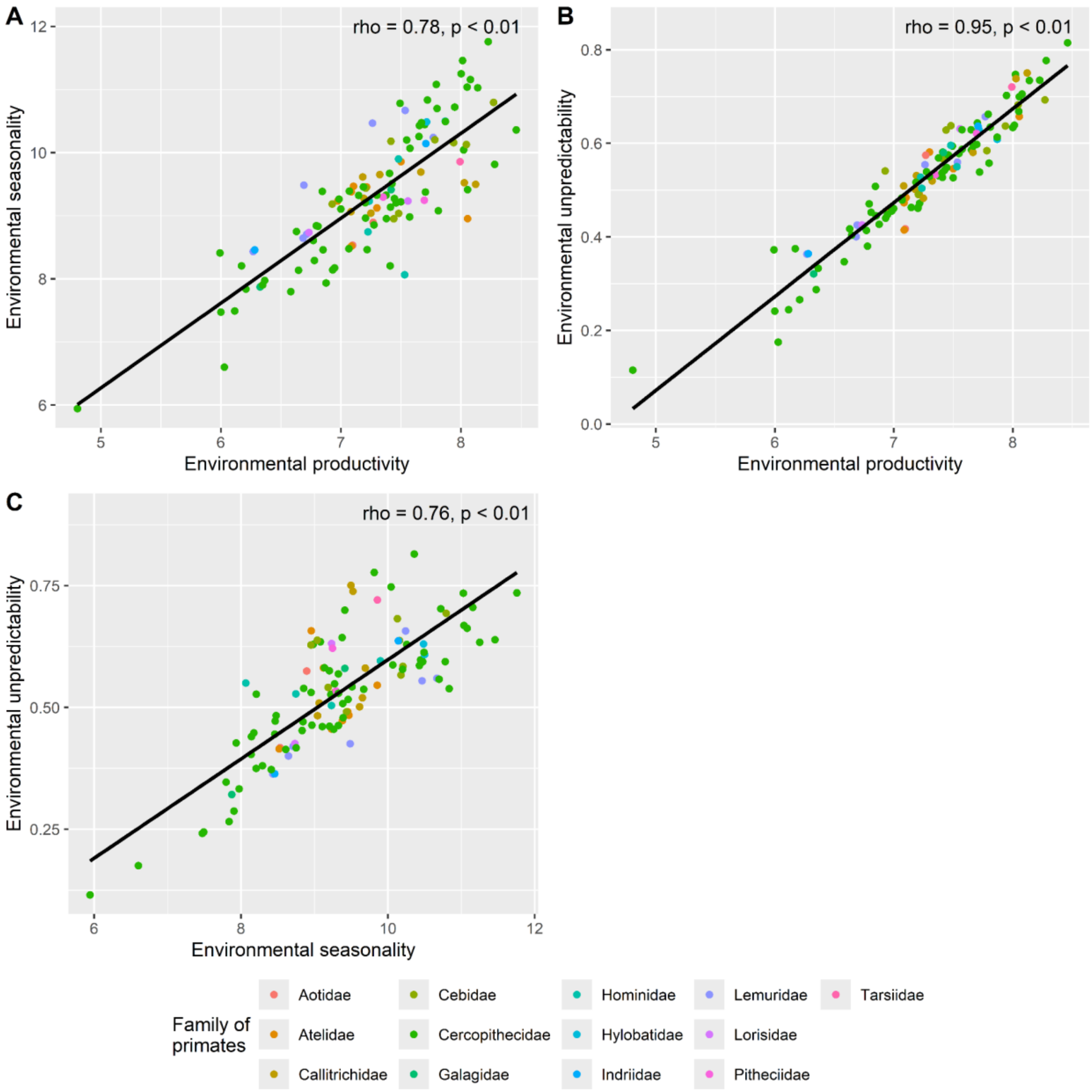

**Figure S3**: Covariation (Spearman's rank correlation *rho* and significance value *p*) between life history traits (diet breadth, allomaternal care and foraging innovations) and environmental productivity (estimated based on monthly rainfall data between 1980 and 2020).

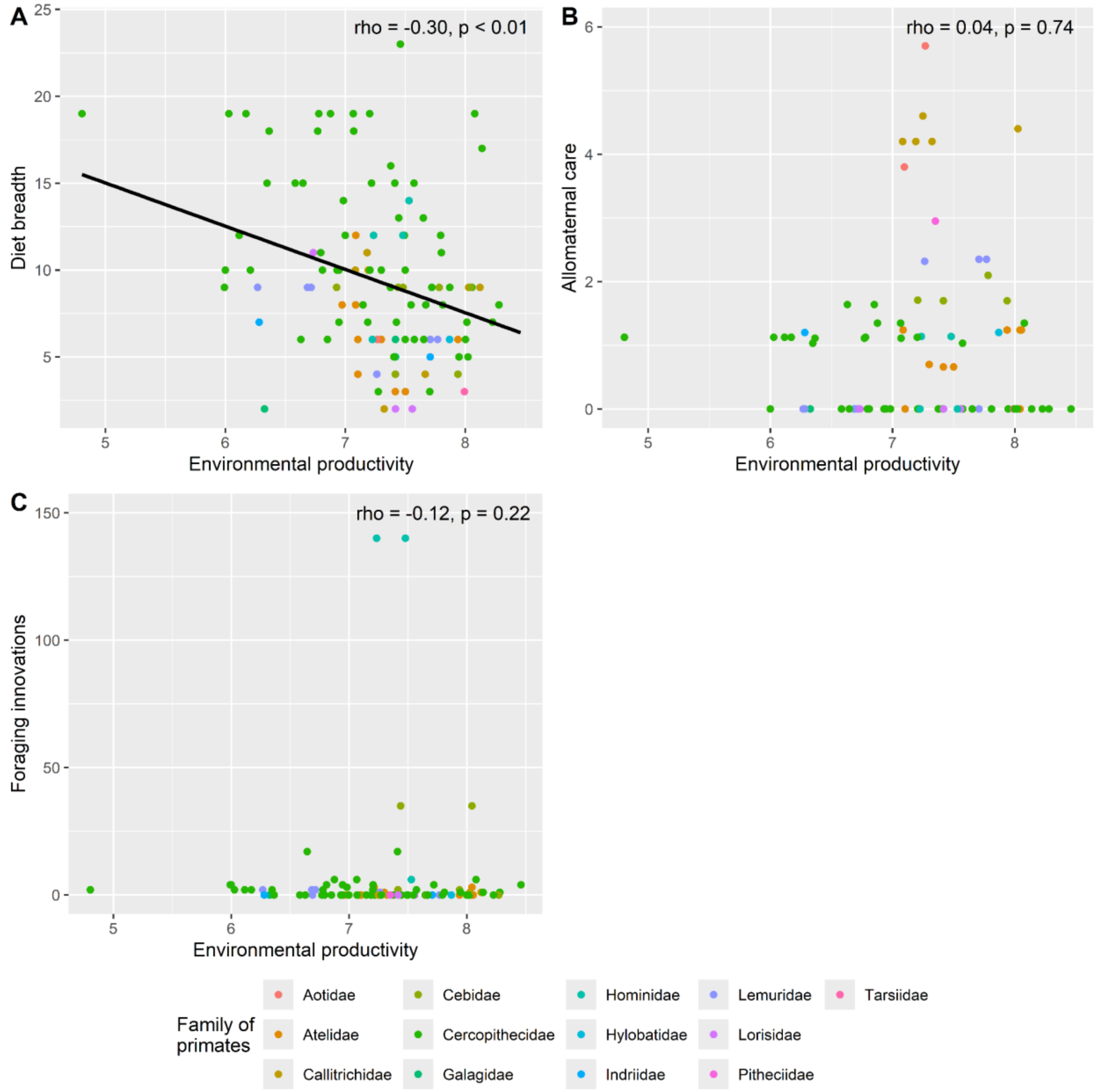

**Figure S4**: Spatial repartition of primate populations from our literature search. We collected *r* values of reproductive seasonality from 132 populations from 94 species of primates.

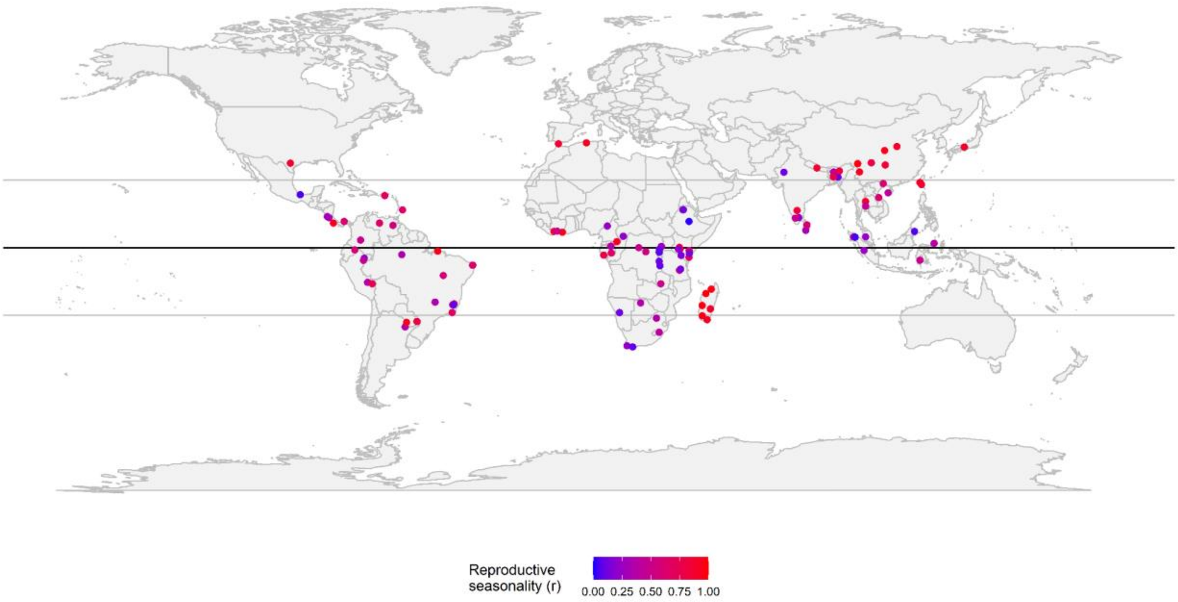

**Figure S5**: Latitudinal range of the populations of primate species ($n$ = 94) included in the analyses.

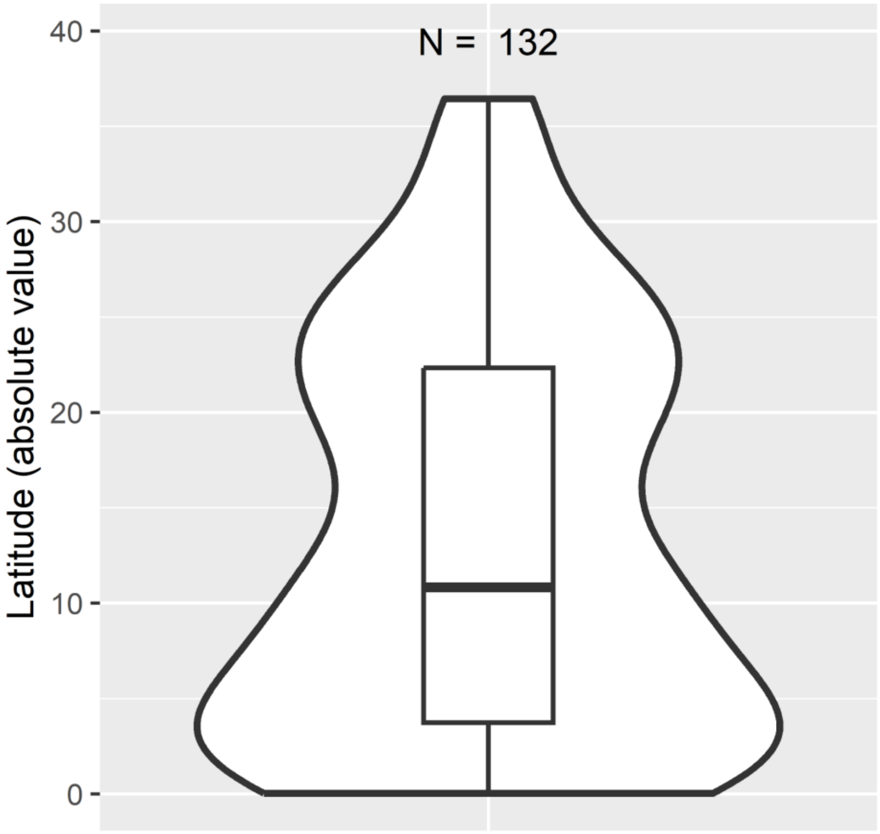

**Figure S6**: Sample bias in relation to taxonomic families of the Primate order. Red bars represent the overall proportion of non-extinct species in each primate family that are represented in the dataset, and blue bars represent the number of populations from each family that are in the dataset.

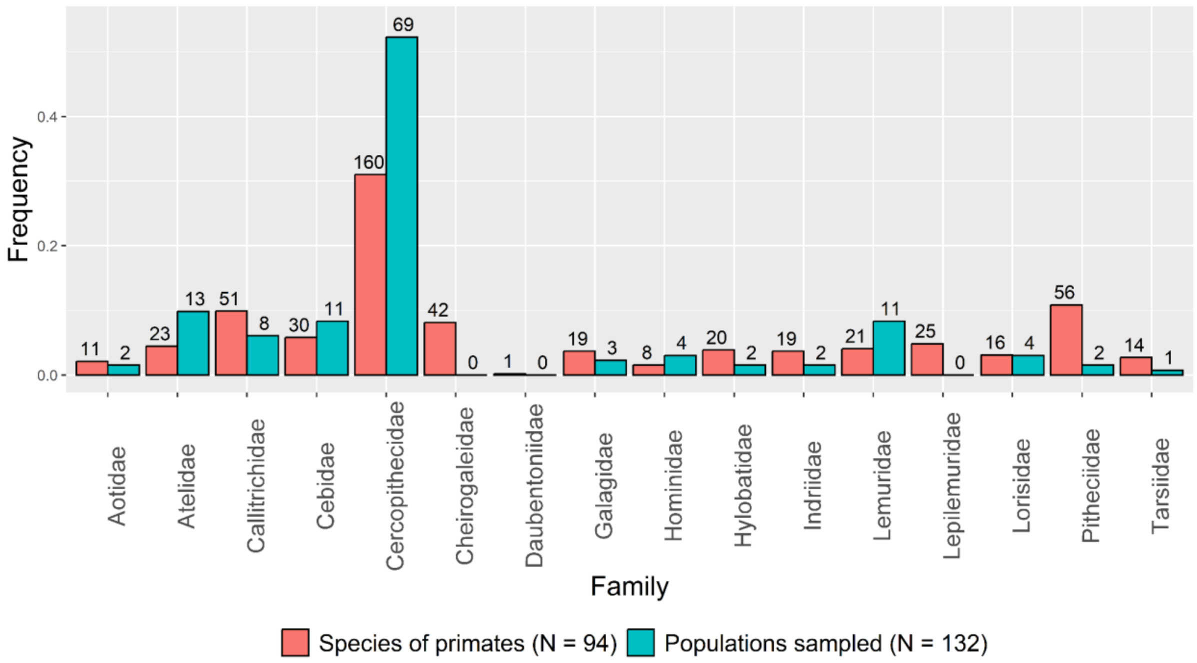

**Figure S7**: Covariation (Spearman's rank correlation *rho* and significance value *p*) of environmental productivity, seasonality and unpredictability (estimated based on monthly rainfall data between 1980 and 2020) with latitude.

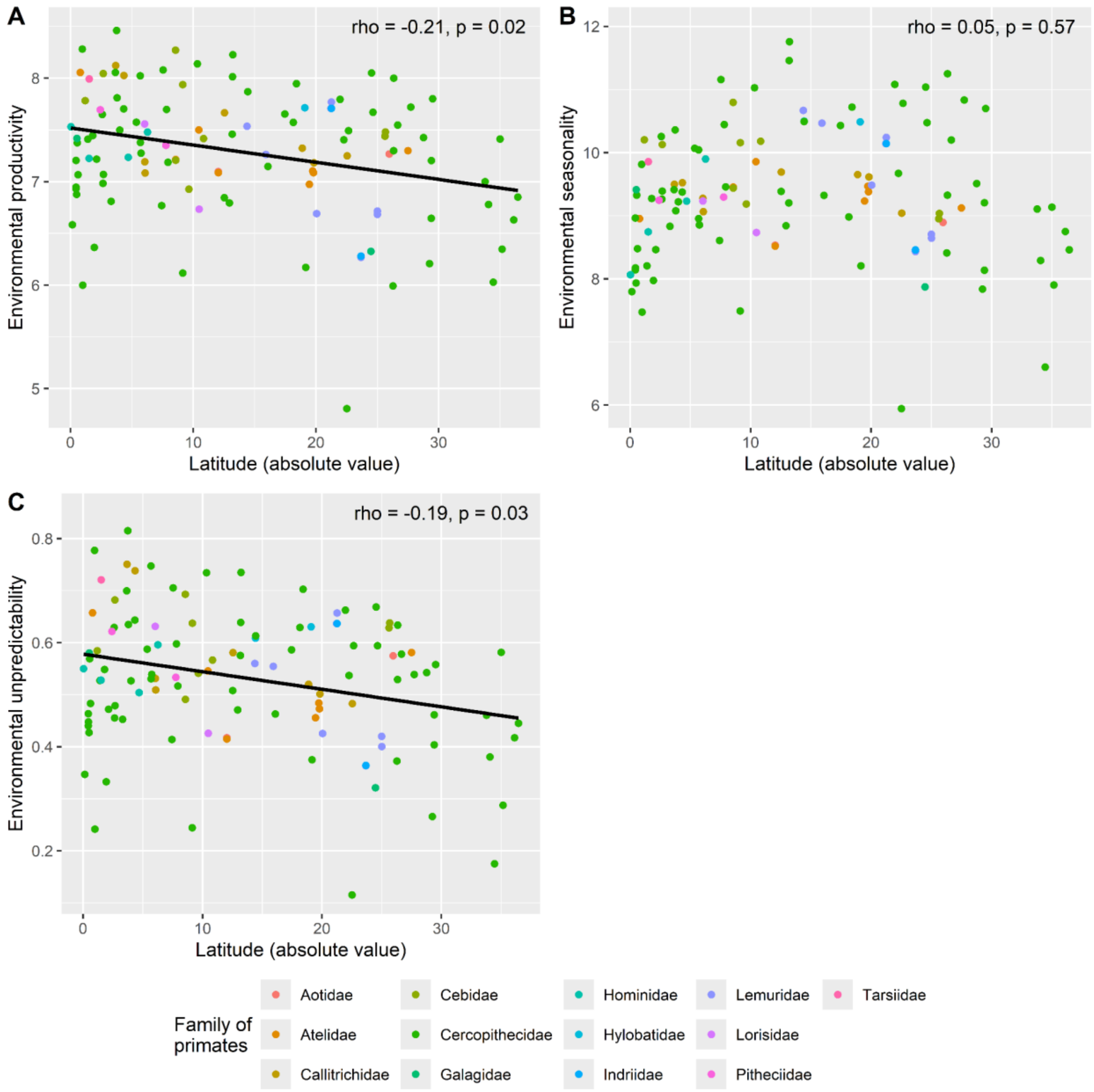

**Figure S8**: Covariation between reproductive seasonality and environmental productivity, seasonality and unpredictability (estimated based on monthly rainfall data between 1980 and 2020). Malagasy species are highlighted with larger dots.

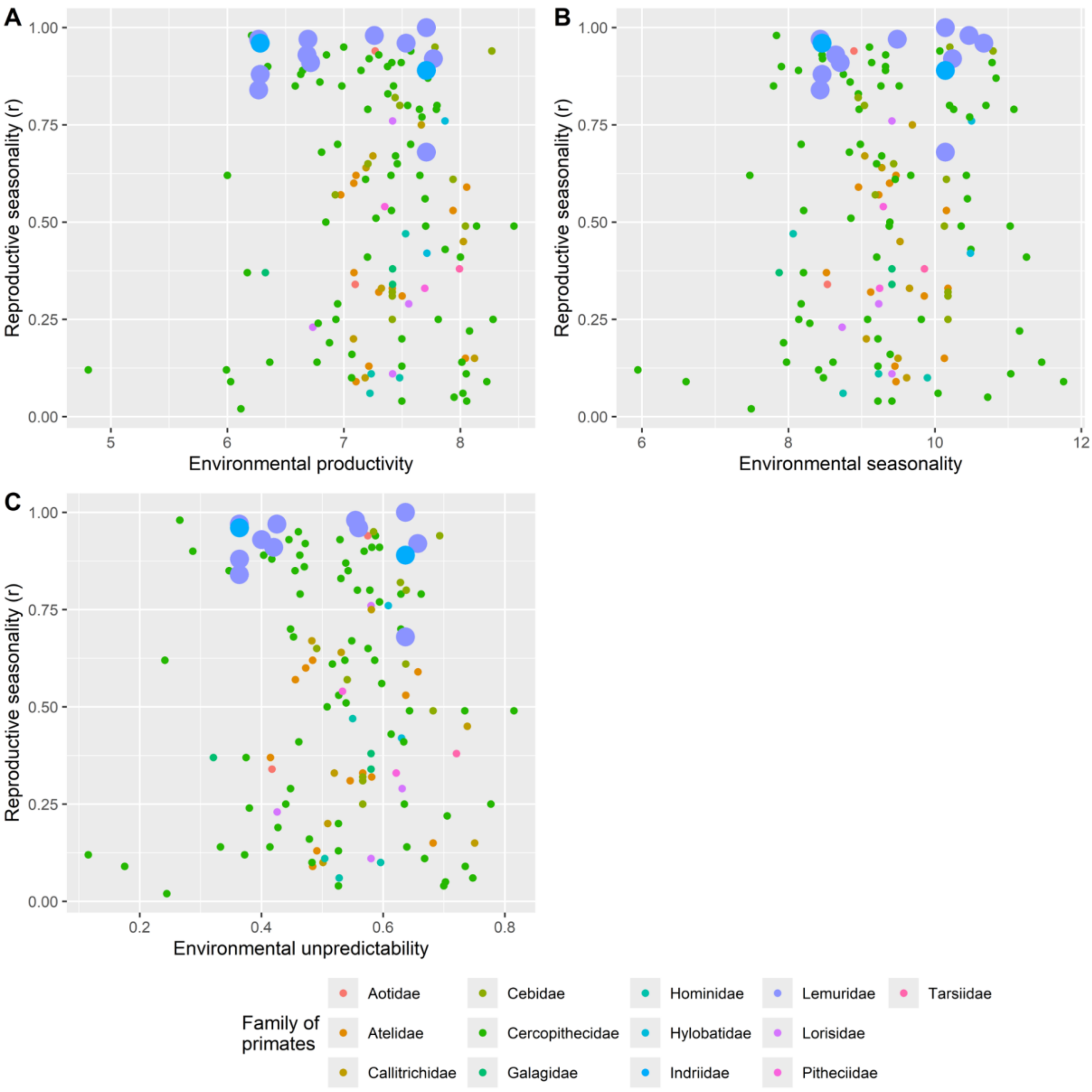

**Table S1**: Dataset used in the comparative analysis investigating reproductive seasonality in primates. Rs: Reproductive seasonality (*r*); Ep: Environmental productivity; Es: Environmental seasonality; Eu: Environmental unpredictability; Ir: Intensity of reproductive effort x 100; Fi: number of Foraging innovations; Ns: Number of studies (foraging innovations); Db: Diet breath; Ac: Allomaternal care; Inf - f: Infanticide by females; Inf - m: Infanticide by males.

| **Family** | **Species** | **Rs** | **Latitude** | **Ep** | **Es** | **Eu** | **Ir** | **Fi** | **Ns** | **Db** | **Ap** | **Inf - f** | **Inf - m** |
|---|---|---|---|---|---|---|---|---|---|---|---|---|---|
| *Aotidae* | *Aotus azarae* | 0.94 | -26.0 | 7.27 | 8.89 | 0.57 | NA | 0 | 22 | 6 | 5.70 | No | No |
| *Aotidae* | *Aotus trivirgatus* | 0.34 | -12.0 | 7.1 | 8.53 | 0.42 | 1.11 | 0 | 58 | NA | 3.80 | NA | NA |
| *Atelidae* | *Alouatta caraya* | 0.32 | -27.5 | 7.3 | 9.12 | 0.58 | NA | 1 | 45 | 6 | 0.70 | No | Yes |
| *Atelidae* | *Alouatta guariba* | 0.09 | -19.7 | 7.1 | 9.47 | 0.48 | NA | 0 | 37 | 4 | 0.00 | No | Yes |
| *Atelidae* | *Alouatta palliata* | 0.31 | 10.5 | 7.5 | 9.86 | 0.55 | 0.27 | 0 | 79 | 3 | 0.66 | No | Yes |
| *Atelidae* | *Alouatta palliata* | 0.33 | 10.8 | 7.42 | 10.18 | 0.57 | 0.27 | 0 | 79 | 3 | 0.66 | No | Yes |
| *Atelidae* | *Alouatta seniculus* | 0.13 | 8.6 | 7.21 | 9.45 | 0.49 | NA | 3 | 82 | NA | 0.00 | No | Yes |
| *Atelidae* | *Alouatta seniculus* | 0.15 | 2.7 | 8.04 | 10.13 | 0.68 | NA | 3 | 82 | NA | 0.00 | No | Yes |
| *Atelidae* | *Ateles belzebuth* | 0.59 | -0.8 | 8.05 | 8.96 | 0.66 | NA | 0 | 12 | 9 | 1.24 | No | Yes |
| *Atelidae* | *Ateles belzebuth* | 0.49 | 2.7 | 8.04 | 10.13 | 0.68 | NA | 0 | 12 | 9 | 1.24 | No | Yes |
| *Atelidae* | *Ateles geoffroyi* | 0.53 | 9.2 | 7.94 | 10.16 | 0.64 | 0.23 | 0 | 58 | 6 | 1.24 | No | Yes |
| *Atelidae* | *Ateles paniscus* | 0.37 | -12.0 | 7.09 | 8.52 | 0.41 | NA | 0 | 30 | 12 | 1.24 | No | No |
| *Atelidae* | *Brachyteles arachnoides* | 0.62 | -19.7 | 7.1 | 9.47 | 0.48 | NA | 0 | 57 | 6 | NA | No | No |

| | | | | | | | | | | | | | |
|---|---|---|---|---|---|---|---|---|---|---|---|---|---|
| *Atelidae* | *Brachyteles hypoxanthus* | 0.60 | -19.8 | 7.09 | 9.38 | 0.47 | NA | NA | NA | 8 | NA | No | No |
| *Atelidae* | *Brachyteles hypoxanthus* | 0.57 | -19.5 | 6.97 | 9.23 | 0.46 | NA | 0 | 57 | 8 | NA | No | No |
| *Callitrichidae* | *Callithrix flaviceps* | 0.10 | -19.8 | 7.18 | 9.61 | 0.5 | NA | NA | NA | 11 | NA | Yes | No |
| *Callitrichidae* | *Callithrix jacchus* | 0.20 | -6.1 | 7.08 | 9.06 | 0.51 | 1.51 | 0 | 161 | 10 | 4.20 | Yes | No |
| *Callitrichidae* | *Callithrix jacchus* | 0.64 | -6.0 | 7.19 | 9.28 | 0.53 | 1.51 | 0 | 161 | 10 | 4.20 | Yes | No |
| *Callitrichidae* | *Callithrix penicillata* | 0.33 | -18.9 | 7.32 | 9.65 | 0.52 | NA | 0 | 0 | 2 | 4.20 | NA | NA |
| *Callitrichidae* | *Cebuella pygmaea* | 0.15 | -3.7 | 8.12 | 9.5 | 0.75 | 1.68 | 1 | 36 | 9 | NA | NA | NA |
| *Callitrichidae* | *Leontopithecus rosalia* | 0.67 | -22.5 | 7.25 | 9.04 | 0.48 | 1.04 | 0 | 85 | NA | 4.60 | No | No |
| *Callitrichidae* | *Saguinus imperator* | 0.75 | -12.6 | 7.66 | 9.69 | 0.58 | NA | 0 | 16 | 4 | NA | NA | NA |
| *Callitrichidae* | *Saguinus mystax* | 0.45 | -4.3 | 8.03 | 9.52 | 0.74 | NA | 1 | 46 | 9 | 4.40 | No | No |
| *Cebidae* | *Cebus capucinus* | 0.32 | 10.8 | 7.42 | 10.18 | 0.57 | 0.52 | 2 | 60 | 4 | 1.70 | No | Yes |
| *Cebidae* | *Cebus capucinus* | 0.61 | 9.2 | 7.94 | 10.16 | 0.64 | 0.52 | 2 | 60 | 4 | 1.70 | No | Yes |
| *Cebidae* | *Cebus capucinus* | 0.25 | 10.8 | 7.42 | 10.18 | 0.57 | NA | NA | NA | 4 | 1.70 | NA | NA |
| *Cebidae* | *Cebus capucinus* | 0.31 | 10.8 | 7.42 | 10.18 | 0.57 | 0.52 | 2 | 60 | 4 | 1.70 | No | Yes |
| *Cebidae* | *Cebus olivaceus* | 0.65 | 8.6 | 7.21 | 9.44 | 0.49 | NA | 2 | 18 | NA | 1.71 | No | Yes |
| *Cebidae* | *Saimiri oerstedii* | 0.94 | 8.6 | 8.27 | 10.8 | 0.69 | NA | 0 | 4 | NA | NA | No | No |
| *Cebidae* | *Saimiri sciureus* | 0.95 | -1.2 | 7.78 | 10.2 | 0.58 | NA | NA | NA | 9 | 2.10 | NA | NA |
| *Cebidae* | *Sapajus apella* | 0.82 | -25.6 | 7.44 | 8.95 | 0.63 | 0.65 | 35 | 249 | 9 | NA | No | Yes |
| *Cebidae* | *Sapajus apella* | 0.49 | 2.7 | 8.04 | 10.13 | 0.68 | 0.65 | 35 | 249 | 9 | NA | No | Yes |

| | | | | | | | | | | | | | |
|---|---|---|---|---|---|---|---|---|---|---|---|---|---|
| *Cebidae* | *Sapajus libidinosus* | 0.57 | -9.7 | 6.93 | 9.19 | 0.54 | NA | NA | NA | 9 | NA | NA | NA |
| *Cebidae* | *Sapajus nigritus* | 0.80 | -25.7 | 7.48 | 9.04 | 0.64 | NA | NA | NA | 9 | NA | No | Yes |
| *Cercopithecidae* | *Cercocebus atys* | 0.83 | 5.7 | 7.38 | 8.95 | 0.53 | NA | NA | NA | 16 | 0.00 | No | Yes |
| *Cercopithecidae* | *Cercocebus sanjei* | 0.56 | -7.8 | 7.7 | 10.44 | 0.6 | 0.67 | NA | NA | NA | NA | NA | NA |
| *Cercopithecidae* | *Cercopithecus ascanius* | 0.25 | 0.4 | 6.93 | 8.14 | 0.44 | NA | 0 | 26 | 10 | 0.00 | No | Yes |
| *Cercopithecidae* | *Cercopithecus campbelli* | 0.94 | 5.4 | 7.57 | 10.07 | 0.59 | NA | NA | NA | 6 | 0.00 | NA | NA |
| *Cercopithecidae* | *Cercopithecus cephus* | 0.04 | 4.0 | 7.5 | 9.22 | 0.53 | NA | 0 | 8 | 10 | NA | NA | NA |
| *Cercopithecidae* | *Cercopithecus lomamiensis* | 0.53 | -1.4 | 7.41 | 8.21 | 0.53 | NA | NA | NA | NA | NA | NA | NA |
| *Cercopithecidae* | *Cercopithecus mitis* | 0.70 | 0.4 | 6.95 | 8.17 | 0.45 | NA | 4 | 56 | 10 | 0.00 | No | Yes |
| *Cercopithecidae* | *Cercopithecus mitis* | 0.68 | -3.3 | 6.81 | 8.83 | 0.45 | NA | 4 | 56 | 10 | 0.00 | No | Yes |
| *Cercopithecidae* | *Cercopithecus mitis* | 0.79 | -0.4 | 7.21 | 8.97 | 0.46 | NA | 4 | 56 | 10 | 0.00 | No | Yes |
| *Cercopithecidae* | *Cercopithecus mitis* | 0.62 | -1.0 | 6 | 7.47 | 0.24 | NA | 4 | 56 | 10 | 0.00 | No | Yes |
| *Cercopithecidae* | *Cercopithecus nictitans* | 0.13 | 4.0 | 7.5 | 9.22 | 0.53 | NA | 0 | 7 | 6 | NA | NA | NA |
| *Cercopithecidae* | *Cercopithecus pogonias* | 0.20 | 4.0 | 7.5 | 9.22 | 0.53 | NA | 0 | 8 | 6 | NA | NA | NA |
| *Cercopithecidae* | *Chlorocebus aethiops* | 0.85 | -2.6 | 6.98 | 9.26 | 0.46 | 0.57 | 3 | 91 | 14 | 0.00 | No | Yes |
| *Cercopithecidae* | *Chlorocebus sabaeus* | 0.65 | 13.2 | 7.46 | 9.2 | 0.58 | 0.52 | NA | NA | 23 | NA | No | NA |
| *Cercopithecidae* | *Erythrocebus patas* | 0.85 | 0.1 | 6.58 | 7.8 | 0.35 | 0.88 | 0 | 33 | 15 | 0.00 | No | Yes |
| *Cercopithecidae* | *Erythrocebus patas* | 0.92 | 2.1 | 7.22 | 8.46 | 0.47 | 0.88 | 0 | 33 | 15 | 0.00 | No | Yes |
| *Cercopithecidae* | *Lophocebus albigena* | 0.29 | 0.4 | 6.95 | 8.17 | 0.45 | 0.79 | 0 | 34 | 7 | NA | No | No |

| | | | | | | | | | | | | | |
|---|---|---|---|---|---|---|---|---|---|---|---|---|---|
| *Cercopithecidae* | *Macaca arctoides* | 0.05 | 18.4 | 7.95 | 10.72 | 0.7 | 0.51 | 1 | 48 | 5 | 0.00 | No | Yes |
| *Cercopithecidae* | *Macaca assamensis* | 0.89 | 16.1 | 7.15 | 9.32 | 0.46 | NA | 0 | 17 | 8 | NA | NA | NA |
| *Cercopithecidae* | *Macaca cyclopis* | 0.79 | 22.0 | 7.79 | 11.08 | 0.66 | NA | 0 | 12 | 12 | NA | NA | NA |
| *Cercopithecidae* | *Macaca cyclopis* | 0.91 | 22.7 | 7.49 | 10.78 | 0.59 | NA | 0 | 12 | 12 | NA | NA | NA |
| *Cercopithecidae* | *Macaca fascicularis* | 0.49 | 3.7 | 8.46 | 10.36 | 0.81 | 0.43 | 4 | 174 | NA | 0.00 | No | Yes |
| *Cercopithecidae* | *Macaca fuscata* | 0.91 | 35.0 | 7.41 | 9.14 | 0.58 | 0.45 | 17 | 253 | 15 | 0.00 | No | Yes |
| *Cercopithecidae* | *Macaca fuscata* | 0.89 | 29.4 | 6.65 | 8.14 | 0.4 | 0.45 | 17 | 253 | 15 | 0.00 | No | Yes |
| *Cercopithecidae* | *Macaca leonina* | 0.11 | 24.5 | 8.05 | 11.04 | 0.67 | 0.45 | NA | NA | 9 | NA | NA | NA |
| *Cercopithecidae* | *Macaca leonina* | 0.43 | 14.4 | 7.87 | 10.49 | 0.61 | 0.45 | NA | NA | 9 | NA | NA | NA |
| *Cercopithecidae* | *Macaca maura* | 0.49 | -4.3 | 7.7 | 9.38 | 0.64 | 0.42 | NA | NA | 3 | NA | NA | NA |
| *Cercopithecidae* | *Macaca mulatta* | 0.70 | 18.2 | 7.57 | 8.98 | 0.63 | 0.47 | 2 | 296 | 15 | 1.03 | Yes | Yes |
| *Cercopithecidae* | *Macaca mulatta* | 0.90 | 35.2 | 6.35 | 7.9 | 0.29 | 0.47 | 2 | 296 | 15 | 1.03 | Yes | Yes |
| *Cercopithecidae* | *Macaca nemestrina* | 0.25 | -1.0 | 8.28 | 9.81 | 0.78 | 0.45 | 1 | 51 | 8 | 0.00 | No | Yes |
| *Cercopithecidae* | *Macaca nemestrina* | 0.25 | 3.8 | 7.81 | 9.08 | 0.63 | 0.45 | 1 | 51 | 8 | 0.00 | No | Yes |
| *Cercopithecidae* | *Macaca radiata* | 0.86 | 12.9 | 6.79 | 8.84 | 0.47 | 0.84 | 0 | 34 | 11 | 0.00 | Yes | Yes |
| *Cercopithecidae* | *Macaca silenus* | 0.49 | 10.3 | 8.14 | 11.03 | 0.73 | NA | 1 | 48 | 17 | 0.00 | No | Yes |
| *Cercopithecidae* | *Macaca sinica* | 0.61 | 7.9 | 7.19 | 9.46 | 0.52 | NA | 0 | 12 | 7 | NA | No | Yes |
| *Cercopithecidae* | *Macaca sylvanus* | 0.93 | 36.5 | 6.85 | 8.46 | 0.45 | 0.97 | 0 | 67 | 6 | 1.64 | Yes | Yes |
| *Cercopithecidae* | *Macaca sylvanus* | 0.88 | 36.1 | 6.63 | 8.75 | 0.42 | 0.97 | 0 | 67 | 6 | 1.64 | Yes | Yes |

| | | | | | | | | | | | | | |
|---|---|---|---|---|---|---|---|---|---|---|---|---|---|
| *Cercopithecidae* | *Macaca thibetana* | 0.80 | 29.5 | 7.8 | 10.7 | 0.56 | 0.4 | NA | NA | 11 | NA | No | Yes |
| *Cercopithecidae* | *Mandrillus sphinx* | 0.67 | -1.8 | 7.44 | 9.27 | 0.55 | 0.52 | 0 | 30 | 13 | NA | No | Yes |
| *Cercopithecidae* | *Mandrillus sphinx* | 0.79 | -2.6 | 7.65 | 10.26 | 0.63 | 0.52 | 0 | 30 | 13 | NA | No | Yes |
| *Cercopithecidae* | *Miopithecus talapoin* | 0.90 | 0.6 | 7.38 | 9.33 | 0.57 | 0.59 | 0 | 4 | 9 | 0.00 | NA | NA |
| *Cercopithecidae* | *Nasalis larvatus* | 0.06 | 5.7 | 8.02 | 10.04 | 0.75 | 0.54 | 0 | 17 | 5 | 0.00 | No | Yes |
| *Cercopithecidae* | *Papio anubis* | 0.19 | -0.5 | 6.88 | 7.93 | 0.43 | 0.57 | 6 | 43 | 19 | 1.35 | No | Yes |
| *Cercopithecidae* | *Papio anubis* | 0.22 | 7.5 | 8.08 | 11.16 | 0.71 | 0.57 | 6 | 43 | 19 | 1.35 | No | Yes |
| *Cercopithecidae* | *Papio anubis* | 0.10 | -0.6 | 7.07 | 8.48 | 0.48 | 0.57 | 6 | 43 | 19 | 1.35 | No | Yes |
| *Cercopithecidae* | *Papio cynocephalus* | 0.14 | -1.9 | 6.36 | 7.98 | 0.33 | 0.59 | 0 | 114 | 18 | 1.11 | Yes | Yes |
| *Cercopithecidae* | *Papio cynocephalus* | 0.14 | -7.4 | 6.77 | 8.61 | 0.41 | 0.59 | 0 | 114 | 18 | 1.11 | Yes | Yes |
| *Cercopithecidae* | *Papio cynocephalus* | 0.16 | -2.7 | 7.07 | 9.39 | 0.48 | 0.59 | 0 | 114 | 18 | 1.11 | Yes | Yes |
| *Cercopithecidae* | *Papio hamadryas* | 0.02 | 9.1 | 6.12 | 7.49 | 0.24 | 0.67 | 2 | 78 | 12 | 1.13 | No | Yes |
| *Cercopithecidae* | *Papio kindae* | 0.50 | -12.5 | 6.84 | 9.39 | 0.51 | NA | NA | NA | NA | NA | No | No |
| *Cercopithecidae* | *Papio ursinus* | 0.24 | -34.1 | 6.78 | 8.29 | 0.38 | 0.55 | 2 | 22 | 19 | 1.13 | Yes | Yes |
| *Cercopithecidae* | *Papio ursinus* | 0.09 | -34.5 | 6.03 | 6.6 | 0.18 | 0.55 | 2 | 22 | 19 | 1.13 | Yes | Yes |
| *Cercopithecidae* | *Papio ursinus* | 0.37 | -19.2 | 6.17 | 8.21 | 0.37 | 0.55 | 2 | 22 | 19 | 1.13 | Yes | Yes |
| *Cercopithecidae* | *Papio ursinus* | 0.41 | -29.4 | 7.2 | 9.21 | 0.46 | 0.55 | 2 | 22 | 19 | 1.13 | Yes | Yes |
| *Cercopithecidae* | *Papio ursinus* | 0.12 | -22.5 | 4.81 | 5.94 | 0.12 | 0.55 | 2 | 22 | 19 | 1.13 | Yes | Yes |
| *Cercopithecidae* | *Presbytis thomasi* | 0.04 | 3.7 | 8.05 | 9.41 | 0.7 | NA | NA | NA | 9 | NA | No | Yes |

| | | | | | | | | | | | | | |
|---|---|---|---|---|---|---|---|---|---|---|---|---|---|
| *Cercopithecidae* | *Procolobus verus* | 0.51 | 5.8 | 7.27 | 8.85 | 0.54 | NA | 0 | 3 | 3 | NA | NA | NA |
| *Cercopithecidae* | *Pygathrix nemaeus* | 0.62 | 17.5 | 7.65 | 10.43 | 0.59 | NA | 0 | 25 | 6 | 0.00 | No | Yes |
| *Cercopithecidae* | *Rhinopithecus bieti* | 0.98 | 29.2 | 6.21 | 7.84 | 0.27 | NA | NA | NA | 10 | NA | No | Yes |
| *Cercopithecidae* | *Rhinopithecus bieti* | 0.93 | 26.3 | 7.3 | 9.33 | 0.53 | NA | NA | NA | 10 | NA | No | Yes |
| *Cercopithecidae* | *Rhinopithecus roxellana* | 0.95 | 33.8 | 7 | 9.11 | 0.46 | NA | 0 | 36 | 12 | NA | No | Yes |
| *Cercopithecidae* | *Semnopithecus entellus* | 0.12 | 26.3 | 5.99 | 8.41 | 0.37 | 0.38 | 4 | 98 | 9 | NA | Yes | Yes |
| *Cercopithecidae* | *Semnopithecus entellus* | 0.87 | 27.7 | 7.72 | 10.83 | 0.54 | 0.38 | 4 | 98 | 9 | NA | Yes | Yes |
| *Cercopithecidae* | *Theropithecus gelada* | 0.09 | 13.2 | 8.23 | 11.76 | 0.74 | 0.54 | 0 | 34 | 7 | 0.00 | No | Yes |
| *Cercopithecidae* | *Theropithecus gelada* | 0.14 | 13.2 | 8.01 | 11.46 | 0.64 | 0.54 | 0 | 34 | 7 | 0.00 | No | Yes |
| *Cercopithecidae* | *Trachypithecus francoisi* | 0.85 | 28.8 | 7.43 | 9.51 | 0.54 | NA | NA | NA | 7 | NA | NA | NA |
| *Cercopithecidae* | *Trachypithecus geei* | 0.41 | 26.3 | 8 | 11.25 | 0.63 | NA | 0 | 7 | 6 | 0.00 | NA | NA |
| *Cercopithecidae* | *Trachypithecus pileatus* | 0.80 | 26.7 | 7.55 | 10.2 | 0.58 | NA | 0 | 5 | 8 | NA | NA | NA |
| *Cercopithecidae* | *Trachypithecus pileatus* | 0.77 | 24.7 | 7.67 | 10.47 | 0.59 | NA | 0 | 5 | 8 | NA | NA | NA |
| *Cercopithecidae* | *Trachypithecus poliocephalus* | 0.62 | 22.3 | 7.4 | 9.67 | 0.54 | NA | NA | NA | 5 | NA | NA | NA |
| *Galagidae* | *Euoticus elegantulus* | 0.38 | 0.5 | 7.42 | 9.41 | 0.58 | NA | 0 | 1 | 5 | NA | NA | NA |
| *Galagidae* | *Galago moholi* | 0.37 | -24.5 | 6.33 | 7.87 | 0.32 | NA | 0 | 14 | 2 | 0.00 | NA | NA |
| *Galagidae* | *Sciurocheirus alleni* | 0.34 | 0.5 | 7.42 | 9.41 | 0.58 | NA | NA | NA | 6 | NA | NA | NA |
| *Hominidae* | *Gorilla beringei* | 0.06 | -1.5 | 7.22 | 8.75 | 0.53 | NA | NA | NA | 6 | 0.00 | No | Yes |
| *Hominidae* | *Pan paniscus* | 0.47 | 0.0 | 7.53 | 8.06 | 0.55 | 0.32 | 6 | 225 | 14 | 0.00 | No | No |

| | | | | | | | | | | | | | |
|---|---|---|---|---|---|---|---|---|---|---|---|---|---|
| *Hominidae* | *Pan troglodytes* | 0.10 | -6.3 | 7.48 | 9.9 | 0.6 | 0.24 | 140 | 755 | 12 | 1.14 | Yes | Yes |
| *Hominidae* | *Pan troglodytes* | 0.11 | -4.7 | 7.23 | 9.23 | 0.5 | 0.24 | 140 | 755 | 12 | 1.14 | Yes | Yes |
| *Hylobatidae* | *Hylobates lar* | 0.76 | 14.4 | 7.87 | 10.5 | 0.61 | 0.21 | 0 | 86 | 6 | 1.20 | No | Yes |
| *Hylobatidae* | *Nomascus hainanus* | 0.42 | 19.1 | 7.71 | 10.49 | 0.63 | NA | NA | NA | NA | NA | NA | NA |
| *Indriidae* | *Propithecus diadema* | 0.89 | -21.3 | 7.71 | 10.14 | 0.64 | NA | 0 | 28 | 5 | NA | No | Yes |
| *Indriidae* | *Propithecus verreauxi* | 0.96 | -23.7 | 6.28 | 8.46 | 0.36 | NA | 0 | 41 | 7 | 1.20 | No | Yes |
| *Lemuridae* | *Eulemur flavifrons* | 0.96 | -14.4 | 7.53 | 10.67 | 0.56 | NA | NA | NA | NA | NA | NA | NA |
| *Lemuridae* | *Eulemur mongoz* | 0.98 | -15.9 | 7.26 | 10.47 | 0.55 | NA | 1 | 13 | 4 | 2.32 | NA | NA |
| *Lemuridae* | *Eulemur rubriventer* | 0.92 | -21.3 | 7.77 | 10.24 | 0.66 | NA | 0 | 13 | 6 | 2.35 | NA | NA |
| *Lemuridae* | *Eulemur rubriventer* | 0.68 | -21.2 | 7.71 | 10.14 | 0.64 | NA | 0 | 13 | 6 | 2.35 | NA | NA |
| *Lemuridae* | *Eulemur rufus* | 0.97 | -20.1 | 6.69 | 9.49 | 0.43 | NA | 0 | 8 | NA | 0.00 | NA | NA |
| *Lemuridae* | *Eulemur rufus* | 1.00 | -21.3 | 7.71 | 10.14 | 0.64 | NA | 0 | 8 | NA | 0.00 | NA | NA |
| *Lemuridae* | *Hapalemur griseus* | 0.88 | -23.7 | 6.28 | 8.46 | 0.36 | NA | 0 | 40 | 7 | 0.00 | No | No |
| *Lemuridae* | *Lemur catta* | 0.91 | -25.0 | 6.71 | 8.71 | 0.42 | NA | 2 | 103 | 9 | 0.00 | Yes | Yes |
| *Lemuridae* | *Lemur catta* | 0.93 | -25.0 | 6.68 | 8.65 | 0.4 | NA | 2 | 103 | 9 | 0.00 | Yes | Yes |
| *Lemuridae* | *Lemur catta* | 0.97 | -23.7 | 6.27 | 8.43 | 0.36 | NA | 2 | 103 | 9 | 0.00 | Yes | Yes |
| *Lemuridae* | *Lemur catta* | 0.84 | -23.7 | 6.27 | 8.43 | 0.36 | NA | 2 | 103 | 9 | 0.00 | Yes | Yes |
| *Lorisidae* | *Arctocebus calabarensis* | 0.11 | 0.5 | 7.42 | 9.41 | 0.58 | 0.96 | 0 | 1 | 4 | NA | NA | NA |
| *Lorisidae* | *Loris lydekkerianus* | 0.23 | 10.5 | 6.73 | 8.73 | 0.43 | NA | NA | NA | 11 | 0.00 | No | No |

| | | | | | | | | | | | | | |
|---|---|---|---|---|---|---|---|---|---|---|---|---|---|
| *Lorisidae* | *Loris tardigradus* | 0.29 | 6.0 | 7.56 | 9.23 | 0.63 | 1.01 | 0 | 14 | 2 | 0.00 | No | No |
| *Lorisidae* | *Perodicticus potto* | 0.76 | 0.5 | 7.42 | 9.41 | 0.58 | NA | 0 | 10 | 2 | 0.00 | NA | NA |
| *Pitheciidae* | *Pithecia chrysocephala* | 0.33 | -2.4 | 7.69 | 9.25 | 0.62 | NA | NA | NA | NA | NA | NA | NA |
| *Pitheciidae* | *Pithecia pithecia* | 0.54 | 7.8 | 7.35 | 9.3 | 0.53 | NA | 0 | 28 | NA | 2.95 | No | No |
| *Tarsiidae* | *Tarsius tarsier* | 0.38 | 1.5 | 7.99 | 9.86 | 0.72 | NA | NA | NA | 3 | 0.00 | NA | NA |

**Table S2**: Reproductive seasonality in primates extracted from the literature. Rs: Reproductive seasonality (*r*); Md: Mean day of birth; Nb: Number of births; Lat: Latitude; Lon: Longitude. These reproductive seasonality values were not computed from raw data, but taken directly from other datasets are indicated with annotations: 1 = Di Bitetti and Janson (2000); 2 = Janson and Verdolin (2005); 3 = Jules Dezeure (personal communication).

| **Family** | **Species** | **Rs** | **Md** | **Nb** | **Location** | **Country** | **Lat** | **Lon** | **Alternative Names** | **References** |
|---|---|---|---|---|---|---|---|---|---|---|
| *Aotidae* | *Aotus azarae* | 0.94 | 304 | 26 | Riacho Pilaga | Argentina | -26.0 | -58.2 | NA | Fernandez-Duque et al. (2002) |
| *Aotidae* | *Aotus trivirgatus* | 0.34 | 325 | NA | Manu | Peru | -12.0 | -71.7 | NA | Janson and Verdolin (2005) |
| *Atelidae* | *Alouatta caraya* | 0.32 | 198 | 74 | Rio Riachuelo | Argentina | -27.5 | -58.7 | NA | Rumiz (1990)[1] |
| *Atelidae* | *Alouatta guariba* | 0.09 | 185 | 34 | Caratinga | Brazil | -19.7 | -41.8 | NA | Strier et al. (2001)[2] |
| *Atelidae* | *Alouatta palliata* | 0.31 | 51 | 27 | Hacienda La Pacifica | Costa Rica | 10.5 | -85.1 | NA | Glander (1980)[1] |
| *Atelidae* | *Alouatta palliata* | 0.33 | 62 | 54 | Santa rosa | Costa Rica | 10.8 | -85.7 | NA | Fedigan et al. (1998)[1] |
| *Atelidae* | *Alouatta seniculus* | 0.13 | 36 | 156 | Hato Masaguaral | Venezuela | 8.6 | -67.6 | NA | Crockett and Rudran (1987)[2] |
| *Atelidae* | *Alouatta seniculus* | 0.15 | 252 | 30 | La Macarena | Colombia | 2.7 | -74.2 | NA | Di Bitetti and Janson (2000) |
| *Atelidae* | *Ateles belzebuth* | 0.59 | 185 | 39 | Yasuni | Ecuador | -0.8 | -76.2 | NA | Ellis et al. (2021) |
| *Atelidae* | *Ateles belzebuth* | 0.49 | 329 | 6 | La Macarena | Colombia | 2.7 | -74.2 | NA | Klein (1971)[2] |

| | | | | | | | | | | |
|---|---|---|---|---|---|---|---|---|---|---|
| *Atelidae* | *Ateles geoffroyi* | 0.53 | 261 | 18 | Barro Colorado Island | Panama | 9.2 | -79.8 | NA | Milton (1981)[1] |
| *Atelidae* | *Ateles paniscus* | 0.37 | 98 | 46 | Manu | Peru | -12 | -71.7 | NA | Di Bitetti and Janson (2000) |
| *Atelidae* | *Brachyteles arachnoides* | 0.62 | 213 | 23 | Caratinga | Brazil | -19.7 | -41.8 | NA | Strier et al. (2001)[2] |
| *Atelidae* | *Brachyteles hypoxanthus* | 0.60 | 205 | 376 | Caratinga | Brazil | -19.8 | -42.1 | NA | Campos et al. (2017) |
| *Atelidae* | *Brachyteles hypoxanthus* | 0.57 | 205 | 28 | Fazenda Monters claros | Brazil | -19.5 | -41.6 | NA | Strier and Ziegler (1994) |
| *Callitrichidae* | *Callithrix flaviceps* | 0.10 | 336 | 10 | Caratinga | Brazil | -19.8 | -41.8 | NA | Ferrari et al. (1996)[1] |
| *Callitrichidae* | *Callithrix jacchus* | 0.20 | 27 | 11 | Rio grande | Brazil | -6.1 | -35.2 | NA | Digby and Barreto (1993)[1] |
| *Callitrichidae* | *Callithrix jacchus* | 0.64 | 30 | 14 | Nisia Floresta | Brazil | -6.0 | -35.2 | NA | Arruda et al. (2005) |
| *Callitrichidae* | *Callithrix penicillata* | 0.33 | 329 | 7 | Mato Gross do Sul | Brazil | -18.9 | -48.3 | NA | Mittermeier et al. (2013) |
| *Callitrichidae* | *Cebuella pygmaea* | 0.15 | 347 | 57 | Pacaya | Peru | -3.7 | -72.8 | NA | Di Bitetti and Janson (2000) |
| *Callitrichidae* | *Leontopithecus rosalia* | 0.67 | 308 | NA | Antas | Brazil | -22.5 | -42.3 | NA | Dietz et al. (1994)[1] |
| *Callitrichidae* | *Saguinus imperator* | 0.75 | 332 | 11 | Los Amigos | Peru | -12.6 | -70.1 | NA | Watsa (2013) |
| *Callitrichidae* | *Saguinus mystax* | 0.45 | 50 | 21 | Quebrada Blanco | Peru | -4.3 | -73.2 | NA | Löttker et al. (2004) |
| *Cebidae* | *Cebus capucinus* | 0.32 | 264 | 40 | Santa rosa | Costa Rica | 10.8 | -85.7 | NA | Di Bitetti and Janson (2000) |

| | | | | | | | | | | |
|---|---|---|---|---|---|---|---|---|---|---|
| *Cebidae* | *Cebus capucinus* | 0.61 | 89 | 25 | Barro Colorado Island | Panama | 9.2 | -79.8 | NA | Di Bitetti and Janson (2000) |
| *Cebidae* | *Cebus capucinus* | 0.25 | 139 | 209 | Santa rosa | Costa Rica | 10.8 | -85.7 | NA | Campos et al. (2017) |
| *Cebidae* | *Cebus capucinus* | 0.31 | 263 | 40 | Santa rosa | Costa Rica | 10.8 | -85.7 | NA | Norconk et al. (1996) |
| *Cebidae* | *Cebus olivaceus* | 0.65 | 172 | 115 | Hato Masaguaral | Venezuela | 8.6 | -67.6 | NA | Robinson (1988)[1] |
| *Cebidae* | *Saimiri oerstedii* | 0.94 | 72 | 37 | Corcovado | Costa Rica | 8.6 | -83.6 | NA | Di Bitetti and Janson (2000) |
| *Cebidae* | *Saimiri sciureus* | 0.95 | 11 | 35 | Ananim | Brazil | -1.2 | -47.3 | NA | Stone and Ruivo (2020) |
| *Cebidae* | *Sapajus apella* | 0.82 | 348 | 36 | Iguazu | Argentina | -25.6 | -54.6 | NA | Di Bitetti and Janson (2000) |
| *Cebidae* | *Sapajus apella* | 0.49 | 89 | 12 | La Macarena | Colombia | 2.7 | -74.2 | NA | Di Bitetti and Janson (2000) |
| *Cebidae* | *Sapajus libidinosus* | 0.57 | 10 | 15 | Fazenda Boa Vista | Brazil | -9.7 | -45.4 | NA | Fragaszy et al. (2016) |
| *Cebidae* | *Sapajus nigritus* | 0.80 | 339 | 40 | Iguazu | Argentina | -25.7 | -54.5 | NA | Di Bitetti and Janson (2001) |
| *Cercopithecidae* | *Cercocebus atys* | 0.83 | 4 | 52 | Tai | Ivory Coast | 5.7 | -6.9 | NA | McGraw et al. (2007) |
| *Cercopithecidae* | *Cercocebus sanjei* | 0.56 | 228 | 28 | Mwanihana forest | United Republic of Tanzania | -7.8 | 36.7 | NA | McCabe and Thompson (2013) |
| *Cercopithecidae* | *Cercopithecus ascanius* | 0.25 | 44 | 7 | Kibale | Uganda | 0.4 | 30.4 | NA | Struhsaker (1977)[2] |
| *Cercopithecidae* | *Cercopithecus campbelli* | 0.94 | 361 | NA | Abidjan | Ivory Coast | 5.4 | -4.0 | NA | Janson and Verdolin (2005) |

| | | | | | | | | | | |
|---|---|---|---|---|---|---|---|---|---|---|
| *Cercopithecidae* | *Cercopithecus cephus* | 0.04 | 28 | 48 | Ngoto forest | Central African Republic | 4.0 | 17.2 | NA | Vanthomme (2010) |
| *Cercopithecidae* | *Cercopithecus lomamiensis* | 0.53 | 250 | 38 | Lomami | Democratic Republic of the Congo | -1.4 | 25.0 | NA | Korchia (2020) |
| *Cercopithecidae* | *Cercopithecus mitis* | 0.70 | 30 | 25 | Kibale | Uganda | 0.4 | 30.4 | *Cercopithecus albogularis; Cercopithecus doggetti; Cercopithecus kandti* | Rudran (1977); Butynski (1982) |
| *Cercopithecidae* | *Cercopithecus mitis* | 0.68 | 57 | 47 | Gede | Kenya | -3.3 | 40.0 | | Foerster et al. (2012) |
| *Cercopithecidae* | *Cercopithecus mitis* | 0.79 | 39 | 26 | Nyeri district | Kenya | -0.4 | 37.0 | | Omar and Vos (1971)[2] |
| *Cercopithecidae* | *Cercopithecus mitis* | 0.62 | 43 | 515 | Kakamega | Kenya | -1.0 | 40.0 | | Campos et al. (2017) |
| *Cercopithecidae* | *Cercopithecus nictitans* | 0.13 | 317 | 18 | Ngoto forest | Central African Republic | 4.0 | 17.2 | NA | Vanthomme (2010) |
| *Cercopithecidae* | *Cercopithecus pogonias* | 0.20 | 269 | 26 | Ngoto forest | Central African Republic | 4.0 | 17.2 | NA | Vanthomme (2010) |
| *Cercopithecidae* | *Chlorocebus aethiops* | 0.85 | 335 | NA | Amboseli | Kenya | -2.6 | 37.2 | NA | Janson and Verdolin (2005) |
| *Cercopithecidae* | *Chlorocebus sabaeus* | 0.65 | 147 | 50 | Barbados | Barbados | 13.2 | -59.6 | NA | Horrocks (1986) |
| *Cercopithecidae* | *Erythrocebus patas* | 0.85 | 2 | NA | Laikipia district | Kenya | 0.1 | 36.7 | NA | Janson and Verdolin (2005) |
| *Cercopithecidae* | *Erythrocebus patas* | 0.92 | 26 | 42 | Kala Maloue | Cameroon | 2.1 | 14.9 | NA | Nakagawa et al. (2003) |
| *Cercopithecidae* | *Lophocebus albigena* | 0.29 | 41 | 76 | Kibale | Uganda | 0.4 | 30.4 | *Lophocebus johnstoni; Lophocebus osmani;* | Arlet et al. (2015) |

| | | | | | | | | | | |
|---|---|---|---|---|---|---|---|---|---|---|
| | | | | | | | | | *Lophocebus ugandae* | |
| *Cercopithecidae* | *Macaca arctoides* | 0.05 | 187 | 24 | Lake Catemaco | Mexico | 18.4 | -95.1 | NA | Estrada and Estrada (1981) |
| *Cercopithecidae* | *Macaca assamensis* | 0.89 | 151 | 29 | Phu Khieo | Thailand | 16.1 | 101.3 | NA | Fürtbauer et al. (2010) |
| *Cercopithecidae* | *Macaca cyclopis* | 0.79 | 121 | 28 | Hengchun | Taiwan | 22.0 | 120.8 | NA | Wu and Lin (1992)[2] |
| *Cercopithecidae* | *Macaca cyclopis* | 0.91 | 128 | 485 | Kaohsiung city | Taiwan | 22.7 | 120.3 | NA | Hsu et al. (2001)[2] |
| *Cercopithecidae* | *Macaca fascicularis* | 0.49 | 257 | 176 | Ketambe | Indonesia | 3.7 | 97.2 | NA | van Noordwijk and van Schaik (1999)[2] |
| *Cercopithecidae* | *Macaca fuscata* | 0.91 | 145 | 416 | Arashiyama | Japan | 35.0 | 135.7 | NA | Fedigan and Griffin (1996) |
| *Cercopithecidae* | *Macaca fuscata* | 0.89 | 140 | 790 | Texas | United States of America | 29.4 | -98.5 | NA | Fedigan and Griffin (1996) |
| *Cercopithecidae* | *Macaca leonina* | 0.11 | 337 | 18 | West Bhanugach Forest | Bangladesh | 24.5 | 91.8 | NA | Feeroz (2003) |
| *Cercopithecidae* | *Macaca leonina* | 0.43 | 104 | 22 | Mo singto | Thailand | 14.4 | 101.4 | NA | Trébouet et al. (2021) |
| *Cercopithecidae* | *Macaca maura* | 0.49 | 127 | 39 | Karaenta | Indonesia | -4.3 | 120.2 | NA | Okamoto et al. (2000) |
| *Cercopithecidae* | *Macaca mulatta* | 0.70 | 34 | 7402 | Cayo santiago | Puerto Rico | 18.2 | -65.7 | NA | Hoffman et al. (2008) |
| *Cercopithecidae* | *Macaca mulatta* | 0.90 | 113 | 64 | Wangwu area | China | 35.2 | 112.3 | NA | Tian et al. (2013) |
| *Cercopithecidae* | *Macaca nemestrina* | 0.25 | 211 | NA | West Sumatra | Indonesia | -1.0 | 100.8 | NA | Janson and Verdolin (2005) |
| *Cercopithecidae* | *Macaca nemestrina* | 0.25 | 262 | NA | Lima Belas | Malaysia | 3.8 | 101.4 | NA | Janson and Verdolin (2005) |
| *Cercopithecidae* | *Macaca radiata* | 0.86 | 68 | 29 | Bangalore | India | 12.9 | 77.6 | NA | Rahaman and Parthasarathy (1969)[2] |

| | | | | | | | | | | |
|---|---|---|---|---|---|---|---|---|---|---|
| *Cercopithecidae* | *Macaca silenus* | 0.49 | 42 | 127 | Anaimalai Hills | India | 10.3 | 77.0 | NA | Sharma et al. (2006) |
| *Cercopithecidae* | *Macaca sinica* | 0.61 | 11 | 50 | Polonnaruwa | Sri Lanka | 7.9 | 81.0 | NA | Janson and Verdolin (2005) |
| *Cercopithecidae* | *Macaca sylvanus* | 0.93 | 136 | 130 | Djurdjura | Algeria | 36.5 | 4.3 | NA | Ménard and Vallet (1993)[2] |
| *Cercopithecidae* | *Macaca sylvanus* | 0.88 | 175 | 147 | Gibraltar | Gibraltar | 36.1 | -5.3 | NA | MacRoberts and MacRoberts (1966)[2] |
| *Cercopithecidae* | *Macaca thibetana* | 0.80 | 73 | 32 | Mont Emei | China | 29.5 | 103.3 | NA | Zhao and Deng (1988)[2] |
| *Cercopithecidae* | *Mandrillus sphinx* | 0.67 | 0 | 218 | Lekedi | Gabon | -1.8 | 13.0 | NA | Jules Dezeure (personal communication) |
| *Cercopithecidae* | *Mandrillus sphinx* | 0.79 | 8 | 208 | Moukalaba Doudou | Gabon | -2.6 | 10.4 | NA | Hongo et al. (2016)[3] |
| *Cercopithecidae* | *Miopithecus talapoin* | 0.90 | 23 | 57 | Makokou | Gabon | 0.6 | 12.8 | *Cercopithecus talapoin* | Gautier-Hion (1970) |
| *Cercopithecidae* | *Nasalis larvatus* | 0.06 | 292 | NA | Kinabatangan | Bornean Malaysia | 5.7 | 118.4 | NA | Boonratana Ramesh (2011) |
| *Cercopithecidae* | *Papio anubis* | 0.19 | 54 | 118 | Gilgil | Kenya | -0.5 | 36.3 | NA | Bercovitch and Harding (1993)[2] |
| *Cercopithecidae* | *Papio anubis* | 0.22 | 352 | 32 | Gashaka Gumti | Nigeria | 7.5 | 11.6 | NA | Higham et al. (2009)[3] |
| *Cercopithecidae* | *Papio anubis* | 0.10 | 18 | 35 | Queen Elisabeth | Uganda | -0.6 | 29.7 | NA | Rowell (1966)[3] |
| *Cercopithecidae* | *Papio cynocephalus* | 0.14 | 330 | 35 | Tana River | Kenya | -1.9 | 40.1 | NA | Bentley-Condit and Smith (1997)[2] |
| *Cercopithecidae* | *Papio cynocephalus* | 0.14 | 203 | 164 | Mikumi | United Republic of Tanzania | -7.4 | 37.1 | NA | Rhine et al. (1988)[3] |
| *Cercopithecidae* | *Papio cynocephalus* | 0.16 | 302 | 820 | Amboseli | Kenya | -2.7 | 37.3 | NA | Campos et al. (2017) |

| | | | | | | | | | | |
|---|---|---|---|---|---|---|---|---|---|---|
| *Cercopithecidae* | *Papio hamadryas* | 0.02 | 166 | 218 | Awash | Ethiopia | 9.1 | 40 | NA | Jules Dezeure (personal communication) |
| *Cercopithecidae* | *Papio kindae* | 0.50 | 197 | 70 | Kasanka | Zambia | -12.5 | 30.2 | NA | Petersdorf et al. (2019)[3] |
| *Cercopithecidae* | *Papio ursinus* | 0.24 | 355 | 52 | Tokai | South Africa | -34.1 | 18.4 | NA | Jules Dezeure (personal communication) |
| *Cercopithecidae* | *Papio ursinus* | 0.09 | 293 | 30 | De Hoop | South Africa | -34.5 | 20.4 | NA | Barrett et al. (2006)[3] |
| *Cercopithecidae* | *Papio ursinus* | 0.37 | 270 | 122 | Moremi | Botswana | -19.2 | 23.2 | NA | Cheney et al. (2004)[3] |
| *Cercopithecidae* | *Papio ursinus* | 0.41 | 327 | 37 | Drakensberg | South Africa | -29.4 | 29.6 | NA | Lycett et al. (1999)[2] |
| *Cercopithecidae* | *Papio ursinus* | 0.12 | 329 | 215 | Tsaobis | Namibia | -22.5 | 15.8 | NA | Jules Dezeure (personal communication) |
| *Cercopithecidae* | *Presbytis thomasi* | 0.04 | 230 | NA | Ketambe | Indonesia | 3.7 | 97.7 | NA | Janson and Verdolin (2005) |
| *Cercopithecidae* | *Procolobus verus* | 0.51 | 345 | 15 | Tai | Ivory Coast | 5.8 | -5.8 | NA | Korstjens and Noe (2004)[2] |
| *Cercopithecidae* | *Pygathrix nemaeus* | 0.62 | 204 | 19 | Hin Namno | Laos | 17.5 | 105.9 | NA | Phiapalath et al. (2011) |
| *Cercopithecidae* | *Rhinopithecus bieti* | 0.98 | 54 | 20 | Honglaxueshan | China | 29.2 | 98.6 | *Pygathrix bieti* | Xiang and Sayers (2009) |
| *Cercopithecidae* | *Rhinopithecus bieti* | 0.93 | 78 | 68 | Yunling | China | 26.3 | 99.2 | | Li et al. (2014) |
| *Cercopithecidae* | *Rhinopithecus roxellana* | 0.95 | 102 | 60 | Zhouzhi | China | 33.8 | 108.0 | *Pygathrix roxellana* | Qi et al. (2008) |
| *Cercopithecidae* | *Semnopithecus entellus* | 0.12 | 100 | 52 | Jodhpur | India | 26.3 | 73.0 | NA | Winkler et al. (1984) |
| *Cercopithecidae* | *Semnopithecus entellus* | 0.87 | 81 | 51 | Ramnagar | Nepal | 27.7 | 84.4 | NA | Koenig et al. (1997)[2] |

| | | | | | | | | | | |
|---|---|---|---|---|---|---|---|---|---|---|
| *Cercopithecidae* | *Theropithecus gelada* | 0.09 | 53 | NA | Sankaber | Ethiopia | 13.2 | 38.0 | NA | Janson and Verdolin (2005) |
| *Cercopithecidae* | *Theropithecus gelada* | 0.14 | 287 | 341 | Simien mountains | Ethiopia | 13.2 | 37.9 | NA | Tinsley Johnson et al. (2018)[3] |
| *Cercopithecidae* | *Trachypithecus francoisi* | 0.85 | 80 | 20 | Mayanghe | China | 28.8 | 108.2 | NA | Wu et al. (2006) |
| *Cercopithecidae* | *Trachypithecus geei* | 0.41 | 188 | 46 | Chakrashila | India | 26.3 | 90.3 | *Presbytis geei* | Shil et al. (2020) |
| *Cercopithecidae* | *Trachypithecus pileatus* | 0.80 | 45 | 17 | Pakhui | India | 26.7 | 92.2 | NA | Solanki et al. (2007) |
| *Cercopithecidae* | *Trachypithecus pileatus* | 0.77 | 68 | NA | Madhupur | Bangladesh | 24.7 | 90.1 | NA | Janson and Verdolin (2005) |
| *Cercopithecidae* | *Trachypithecus poliocephalus* | 0.62 | 29 | 133 | Nongguan Karst | China | 22.3 | 107.5 | NA | Jin et al. (2009) |
| *Galagidae* | *Euoticus elegantulus* | 0.38 | 24 | NA | Makokou | Gabon | 0.5 | 12.8 | NA | Charles-Dominique (1977)[2] |
| *Galagidae* | *Galago moholi* | 0.37 | 344 | 4 | Nylsvley | South Africa | -24.5 | 28.7 | NA | Pullen et al. (2000)[2] |
| *Galagidae* | *Sciurocheirus alleni* | 0.34 | 22 | NA | Makokou | Gabon | 0.5 | 12.8 | NA | Charles-Dominique (1977) |
| *Hominidae* | *Gorilla beringei* | 0.06 | 180 | 269 | Karisoke | Rwanda | -1.5 | 29.6 | NA | Campos et al. (2017) |
| *Hominidae* | *Pan paniscus* | 0.47 | 68 | 15 | Wamba | Democratic Republic of the Congo | 0.0 | 22.6 | NA | Furuichi et al. (1998) |
| *Hominidae* | *Pan troglodytes* | 0.10 | 71 | 85 | Mahale mountain | United Republic of Tanzania | -6.3 | 29.9 | NA | Nishida et al. (1990)[2] |
| *Hominidae* | *Pan troglodytes* | 0.11 | 197 | 198 | Gombe | United Republic of Tanzania | -4.7 | 29.6 | NA | Campos et al. (2017) |

| | | | | | | | | | | |
|---|---|---|---|---|---|---|---|---|---|---|
| *Hylobatidae* | *Hylobates lar* | 0.76 | NA | 15 | Khao Yai | Thailand | 14.4 | 101.4 | NA | Savini et al. (2008) |
| *Hylobatidae* | *Nomascus hainanus* | 0.42 | 15 | 17 | Bawangling | China | 19.1 | 109.2 | NA | Deng et al. (2017) |
| *Indriidae* | *Propithecus diadema* | 0.89 | 158 | NA | Ranomafana | Madagascar | -21.3 | 47.4 | NA | Janson and Verdolin (2005) |
| *Indriidae* | *Propithecus verreauxi* | 0.96 | 204 | 706 | Beza Mahafaly | Madagascar | -23.7 | 44.6 | NA | Campos et al. (2017) |
| *Lemuridae* | *Eulemur flavifrons* | 0.96 | 254 | 22 | Sahamalaza | Madagascar | -14.4 | 47.8 | NA | Volampeno et al. (2011) |
| *Lemuridae* | *Eulemur mongoz* | 0.98 | 293 | 6 | Anjamena | Madagascar | -15.9 | 45.8 | NA | Curtis and Zaramody (1999)[2] |
| *Lemuridae* | *Eulemur rubriventer* | 0.92 | 282 | NA | Ranomafana | Madagascar | -21.3 | 47.4 | NA | Janson and Verdolin (2005) |
| *Lemuridae* | *Eulemur rubriventer* | 0.68 | 286 | 30 | Ranomafana | Madagascar | -21.2 | 47.5 | NA | Tecot (2010) |
| *Lemuridae* | *Eulemur rufus* | 0.97 | 282 | 45 | Kirindi Forest | Madagascar | -20.1 | 44.5 | NA | Ostner and Kappeler (2004) |
| *Lemuridae* | *Eulemur rufus* | 1.00 | 262 | NA | Ranomafana | Madagascar | -21.3 | 47.4 | NA | Janson and Verdolin (2005) |
| *Lemuridae* | *Hapalemur griseus* | 0.88 | 334 | NA | Beza Mahafaly | Madagascar | -23.7 | 44.6 | NA | Janson and Verdolin (2005) |
| *Lemuridae* | *Lemur catta* | 0.91 | 259 | 178 | Berenty | Madagascar | -25.0 | 46.3 | NA | Koyama et al. (2001) |
| *Lemuridae* | *Lemur catta* | 0.93 | 257 | 44 | Berenty | Madagascar | -25.0 | 46.3 | NA | Jones (1983) |
| *Lemuridae* | *Lemur catta* | 0.97 | 251 | NA | Beza | Madagascar | -23.7 | 44.6 | NA | Janson and Verdolin (2005) |
| *Lemuridae* | *Lemur catta* | 0.84 | 94 | 103 | Beza | Madagascar | -23.7 | 44.6 | NA | Pereira (1991) |

| | | | | | | | | | | |
|---|---|---|---|---|---|---|---|---|---|---|
| *Lorisidae* | *Arctocebus calabarensis* | 0.11 | 17 | NA | Makokou | Gabon | 0.5 | 12.8 | NA | Charles-Dominique (1977)[2] |
| *Lorisidae* | *Loris lydekkerianus* | 0.23 | 106 | 14 | Dindigul | India | 10.5 | 78.2 | NA | Radhakrishna and Singh (2004) |
| *Lorisidae* | *Loris tardigradus* | 0.29 | 76 | 8 | Masmullah Proposed | Sri Lanka | 6.0 | 80.6 | NA | Nekaris (2003) |
| *Lorisidae* | *Perodicticus potto* | 0.76 | 272 | NA | Makokou | Gabon | 0.5 | 12.8 | NA | Charles-Dominique (1977)[2] |
| *Pitheciidae* | *Pithecia chrysocephala* | 0.33 | 364 | 6 | Colosso | Brazil | -2.4 | -59.8 | NA | Setz and Gaspar (1997) |
| *Pitheciidae* | *Pithecia pithecia* | 0.54 | 16 | 10 | Isla Redonda | Venezuela | 7.8 | -62.9 | NA | Norconk (2006) |
| *Tarsiidae* | *Tarsius tarsier* | 0.38 | 113 | 43 | Tangkoko | Indonesia | 1.5 | 125.2 | NA | Gursky (2007) |

**References supplementary material**

Arlet, M.E., Isbell, L.A., Kaasik, A., Molleman, F., Chancellor, R.L. and Chapman, C.A. et al. (2015) Determinants of Reproductive Performance Among Female Gray-Cheeked Mangabeys (*Lophocebus albigena*) in Kibale National Park, Uganda. International Journal of Primatology, 36(1), 55–73. Available from: https://doi.org/10.1007/s10764-014-9810-4

Arruda, M.F., Araújo, A., Sousa, M.B.C., Albuquerque, F.S., Albuquerque, A.C.S.R. and Yamamoto, M.E. (2005) Two breeding females within free-living groups may not always indicate polygyny: alternative subordinate female strategies in common marmosets (*Callithrix jacchus*). Folia Primatologica; International Journal of Primatology, 76(1), 10–20. Available from: https://doi.org/10.1159/000082451

Barrett, L., Peter Henzi, S. and Lycett, J.E. (2006) Whose Life Is It Anyway? Maternal Investment, Developmental Trajectories, and Life History Strategies in Baboons. In: Swedell, L. and Leigh, S.R. (Eds.) Reproduction and Fitness in Baboons: Behavioral, Ecological, and Life History Perspectives. Springer US: New York, NY, pp. 199–224.

Bentley-Condit, V.K. and Smith, E.O. (1997) Female Reproductive Parameters of Tana River Yellow Baboons. International Journal of Primatology, 18(4), 581–596. Available from: https://doi.org/10.1023/A:1026315323400

Bercovitch, F.B. and Harding, R.S. (1993) Annual birth patterns of savanna baboons (*Papio cynocephalus anubis*) over a ten-year period at Gilgil, Kenya. Folia Primatologica; International Journal of Primatology, 61(3), 115–122. Available from: https://doi.org/10.1159/000156738

Boonratana Ramesh (2011) Observations on the sexual behavior and birth seasonality of proposcis monkey (*Nasalis larvatus*) along the lower Kinabatangan River, northern Borneo. Asian Primates Journal, 2(1), 2–9.

Butynski, T.M. (1982) Harem-male replacement and infanticide in the blue monkey (*Cercopithecus mitus stuhlmanni*) in the Kibale Forest, Uganda. American Journal of Primatology, 3(1-4), 1–22. Available from: https://doi.org/10.1002/ajp.1350030102

Campos, F.A., Morris, W.F., Alberts, S.C., Altmann, J., Brockman, D.K. and Cords, M. et al. (2017) Does climate variability influence the demography of wild primates? Evidence from longterm life-history data in seven species. Global Change Biology, 23(11), 4907–4921. Available from: https://doi.org/10.1111/gcb.13754

Charles-Dominique, P. (1977) Ecology and behaviour of nocturnal primates: Prosimians of equatorial West Africa. Columbia University Press: New York.

Cheney, D.L., Seyfarth, R.M., Fischer, J., Beehner, J., Bergman, T. and Johnson, S.E. et al. (2004) Factors Affecting Reproduction and Mortality Among Baboons in the Okavango Delta, Botswana. International Journal of Primatology, 25(2), 401–428. Available from: https://doi.org/10.1023/B:IJOP.0000019159.75573.13

Crockett, C.M. and Rudran, R. (1987) Red howler monkey birth data I: Seasonal variation. American Journal of Primatology, 13(4), 347–368. Available from: https://doi.org/10.1002/ajp.1350130402

Curtis, D.J. and Zaramody, A. (1999) Social structure and seasonal variation in the behaviour of *Eulemur mongoz*. Folia Primatologica; International Journal of Primatology, 70(2), 79–96. Available from: https://doi.org/10.1159/000021679

Deng, H., Zhang, M. and Zhou, J. (2017) Recovery of the Critically Endangered Hainan gibbon *Nomascus hainanus*. Oryx, 51(1), 161–165. Available from: https://doi.org/10.1017/S0030605315000678

Di Bitetti, M.S. and Janson, C.H. (2000) When will the stork arrive? Patterns of birth seasonality in neotropical primates. American journal of primatology, 50(2), 109–130. Available from: https://doi.org/10.1002/(SICI)1098-2345(200002)50:2<109::AID-AJP2>3.0.CO;2-W

Di Bitetti, M.S. and Janson, C.H. (2001) Reproductive Socioecology of Tufted Capuchins (*Cebus apella nigritus*) in Northeastern Argentina. International Journal of Primatology, 22(2), 127–142. Available from: https://doi.org/10.1023/A:1005611228927

Dietz, J.M., Baker, A.J. and Miglioretti, D. (1994) Seasonal variation in reproduction, juvenile growth, and adult body mass in golden lion tamarins (*Leontopithecus rosalia*). American Journal of Primatology, 34(2), 115–132. Available from: https://doi.org/10.1002/ajp.1350340204

Digby, L.J. and Barreto, C.E. (1993) Social organization in a wild population of *Callithrix jacchus*: Group Composition and Dynamics. Folia Primatologica; International Journal of Primatology, 61(3), 123–134. Available from: https://doi.org/10.1159/000156739

Ellis, K.M., Abondano, L.A., Montes-Rojas, A., Link, A. and Di Fiore, A. (2021) Reproductive seasonality in two sympatric primates (*Ateles belzebuth* and *Lagothrix lagotricha poeppigii*) from Amazonian Ecuador. American Journal of Primatology, 83(1), e23220. Available from: https://doi.org/10.1002/ajp.23220

Estrada, A. and Estrada, R. (1981) Reproductive seasonality in a free-ranging troop of stumptail macaques (*Macaca arctoides*): A five-year report. Primates, 22(4), 503–511. Available from: https://doi.org/10.1007/BF02381242

Fedigan, L.M. and Griffin, L. (1996) Determinants of reproductive seasonality in the Arashiyama West Japanese macaques. In: Fa, J.E. and Lindburg, D.G. (Eds.)

Evolution and ecology of macaque societies. Cambridge University Press: Cambridge.

Fedigan, L.M., Rose, L.M. and Avila, R.M. (1998) Growth of Mantled Howler Groups in a Regenerating Costa Rican Dry Forest. International Journal of Primatology, 19(3), 405–432. Available from: https://doi.org/10.1023/A:1020304304558

Feeroz, M.M. (2003) Breeding activities of the Pig-tailed Macaque (*Macaca leonina*) in Bangladesh. Zoos' Print Journal, 18(8), 1175–1179. Available from: https://doi.org/10.11609/JoTT.ZPJ.18.8.1175-9

Fernandez-Duque, E., Rotundo, M. and Ramirez-Llorens, P. (2002) Environmental Determinants of Birth Seasonality in Night Monkeys (*Aotus azarai*) of the Argentinean Chaco. International Journal of Primatology, 23(3), 639–656. Available from: https://doi.org/10.1023/A:1014929902923

Ferrari, S.F., Kátia, H., Corrêa, M. and Coutinho, P.E.G. (1996) Ecology of the "Southern" Marmosets (*Callithrix aurita* and *Callithrix flaviceps*). In: Norconk, M.A., Rosenberger, A.L. and Garber, P.A. (Eds.) Adaptive Radiations of Neotropical Primates. Springer Science & Business Media, pp. 157–171.

Foerster, S., Cords, M. and Monfort, S.L. (2012) Seasonal energetic stress in a tropical forest primate: proximate causes and evolutionary implications. PloS One, 7(11), e50108. Available from: https://doi.org/10.1371/journal.pone.0050108

Fragaszy, D.M., Izar, P., Liu, Q., Eshchar, Y., Young, L.A. and Visalberghi, E. (2016) Body mass in wild bearded capuchins, (*Sapajus libidinosus*): Ontogeny and sexual dimorphism. American Journal of Primatology, 78(4), 473–484. Available from: https://doi.org/10.1002/ajp.22509

Fürtbauer, I., Schülke, O., Heistermann, M. and Ostner, J. (2010) Reproductive and Life History Parameters of Wild Female *Macaca assamensis*. International Journal of

Primatology, 31(4), 501–517. Available from: https://doi.org/10.1007/s10764-010-9409-3

Furuichi, T., Idani, G., Ihobe, H., Kuroda, S., Kitamura, K. and Mori, A. et al. (1998) Population Dynamics of Wild Bonobos (*Pan paniscus*) at Wamba. International Journal of Primatology, 19(6), 1029–1043. Available from: https://doi.org/10.1023/A:1020326304074

Gautier-Hion, A. (1970) L'organisation sociale d'une bande de Talapoins (*Miopithecus talapoin*) dans le nord-est du Gabon. Folia primatologica; international journal of primatology, 12(2), 116–141. Available from: https://doi.org/10.1159/000155285

Glander, K.E. (1980) Reproduction and population growth in free-ranging mantled howling monkeys. American Journal of Physical Anthropology, 53(1), 25–36. Available from: https://doi.org/10.1002/ajpa.1330530106

Gursky, S. (2007) The spectral tarsier. Pearson/Prentice Hall: Upper Saddle River.

Heldstab, S.A., van Schaik, C.P., Müller, D.W.H., Rensch, E., Lackey, L.B. and Zerbe, P. et al. (2021) Reproductive seasonality in primates: patterns, concepts and unsolved questions. Biological Reviews of the Cambridge Philosophical Society, 96(1), 66–88. Available from: https://doi.org/10.1111/brv.12646

Higham, J.P., Warren, Y., Adanu, J., Umaru, B.N., MacLarnon, A.M. and Sommer, V. et al. (2009) Living on the edge: life-history of olive baboons at Gashaka-Gumti National Park, Nigeria. American Journal of Primatology, 71(4), 293–304. Available from: https://doi.org/10.1002/ajp.20651

Hoffman, C.L., Ruiz-Lambides, A.V., Davila, E., Maldonado, E., Gerald, M.S. and Maestripieri, D. (2008) Sex differences in survival costs of reproduction in a promiscuous primate. Behavioral Ecology and Sociobiology, 62(11), 1711–1718. Available from: https://doi.org/10.1007/s00265-008-0599-z

Hongo, S., Nakashima, Y., Akomo-Okoue, E.F. and Mindonga-Nguelet, F.L. (2016) Female Reproductive Seasonality and Male Influxes in Wild Mandrills (*Mandrillus sphinx*). International Journal of Primatology, 37(3), 416–437. Available from: https://doi.org/10.1007/s10764-016-9909-x

Horrocks, J.A. (1986) Life-history characteristics of a wild population of vervets (*Cercopithecus aethiops sabaeus*) in Barbados, West Indies. International Journal of Primatology, 7(1), 31–47. Available from: https://doi.org/10.1007/BF02692308

Hsu, M.J., Agoramoorthy, G. and Lin, J.-F. (2001) Birth seasonality and interbirth intervals in freeranging formosan macaques,*Macaca cyclopis*, at Mt. Longevity, Taiwan. Primates, 42(1), 15–25. Available from: https://doi.org/10.1007/BF02640685

Janson, C. and Verdolin, J. (2005) Seasonality of primate births in relation to climate. In: Brockman, D.K. and van Schaik, C. (Eds.) Seasonality in primates: Implications for human evolution. Cambridge University Press: Cambridge, pp. 307–350.

Jin, T., Wang, D.-Z., Zhao, Q., Yin, L.-J., Qin, D.-G. and Ran, W.-Z. et al. (2009) Reproductive parameters of wild *Trachypithecus leucocephalus*: seasonality, infant mortality and interbirth interval. American Journal of Primatology, 71(7), 558–566. Available from: https://doi.org/10.1002/ajp.20688

Jones, K.C. (1983) Inter-troop transfer of *Lemur catta* males at Berenty, Madagascar. Folia Primatologica; International Journal of Primatology, 40(1-2), 145–160. Available from: https://doi.org/10.1159/000156096

Klein, L.L. (1971) Observations on Copulation and Seasonal Reproduction of Two Species of Spider Monkeys, *Ateles belzebuth* and *A. geoffroyi*. Folia primatologica; international journal of primatology, 15(3-4), 233–248. Available from: https://doi.org/10.1159/000155382

Koenig, A., Borries, C., Chalise, M.K. and Winkler, P. (1997) Ecology, nutrition, and timing of reproductive events in an Asian primate, the Hanuman langur (*Presbytis entellus*). Journal of Zoology, 243(2), 215–235. Available from: https://doi.org/10.1111/j.1469-7998.1997.tb02778.x

Korchia, C.S.F. (2020) Behavioral ecology of *Cercopithecus lomamiensis* in the Lomami. Florida Atlantic University.

Korstjens, A.H. and Noë, R. (2004) Mating system of an exceptional primate, the olive colobus (*Procolobus verus*). American Journal of Primatology, 62(4), 261–273. Available from: https://doi.org/10.1002/ajp.20020

Koyama, N., Nakamichi, M., Oda, R., Miyamoto, N., Ichino, S. and Takahata, Y. (2001) A ten-year summary of reproductive parameters for ring-tailed lemurs at berenty, Madagascar. Primates, 42(1), 1–14. Available from: https://doi.org/10.1007/BF02640684

Li, J.-F., He, Y.-C., Huang, Z.-P., Wang, S.-J., Xiang, Z.-F. and Zhao, J.-J. et al. (2014) Birth seasonality and pattern in black-and-white snub-nosed monkeys (*Rhinopithecus bieti*) at Mt. Lasha, Yunnan. Dong Wu Xue Yan Jiu = Zoological Research, 35(6), 474–484. Available from: https://doi.org/10.13918/j.issn.2095-8137.2014.6.474

Löttker, P., Huck, M. and Heymann, E.W. (2004) Demographic parameters and events in wild moustached Tamarins (*Saguinus mystax*). American Journal of Primatology, 64(4), 425–449. Available from: https://doi.org/10.1002/ajp.20090

Lycett, J.E., Weingrill, T. and Henzia, S.P. (1999) Birth patterns in the Drakensberg Mountain baboons (*Papio cynocephalus ursinus*). South African Journal of Science, 95(8), 354–356. Available from: https://hdl.handle.net/10520/AJA00382353_8360

Machado, F. F., Jardim, L., Dinnage, R., Brito, D. and Cardillo, M. (2022) Diet disparity and diversity predict extinction risk in primates. Animal Conservation, 26(3), 331-339. Available from: https://doi.org/10.1111/acv.12823

MacRoberts, M.H. and MacRoberts, B.R. (1966) The annual reproductive cycle of the Barbary Ape (*Macaca sylvana*) in Gibraltar. American Journal of Physical Anthropology, 25(3), 299–304. Available from: https://doi.org/10.1002/ajpa.1330250309

McCabe, G.M. and Thompson, M.E. (2013) Reproductive seasonality in wild Sanje mangabeys (*Cercocebus sanjei*), Tanzania: Relationship between the capital breeding strategy and infant survival. Behaviour, 150(12), 1399–1429. Available from: https://doi.org/10.1163/1568539X-00003102

McGraw, W.S., Zuberbühler, K. and Noë, R. (Eds.) (2007) The Monkeys of the Taï Forest: An African primate community. Cambridge University Press: Cambridge, New York.

Ménard, N. and Vallet, D. (1993) Population dynamics of *Macaca sylvanus* in Algeria: An 8-year study. American Journal of Primatology, 30(2), 101–118. Available from: https://doi.org/10.1002/ajp.1350300203

Milton, K. (1981) Estimates of reproductive parameters for free-ranging *Ateles geoffroyi*. Primates, 22(4), 574–579. Available from: https://doi.org/10.1007/bf02381250

Mittermeier, R.A., Wilson, D.E., Rylands, A.B., Martínez-Vilalta, A., Leslie, D.M. and Richardson, M. (Eds.) (2013) Handbook of the mammals of the world: vol. 3: Primates. Lynx Edicions: Barcelona.

Nakagawa, N., Ohsawa, H. and Muroyama, Y. (2003) Life-history parameters of a wild group of West African patas monkeys (*Erythrocebus patas patas*). Primates; Journal of Primatology, 44(3), 281–290. Available from: https://doi.org/10.1007/s10329-003-0042-z

Nekaris, K.A.I. (2003) Observations of mating, birthing and parental behaviour in three subspecies of slender loris (*Loris tardigradus* and *Loris lydekkerianus*) in India and Sri Lanka. Folia Primatologica; International Journal of Primatology, 74(5-6), 312–336. Available from: https://doi.org/10.1159/000073317

Nishida, T., Takasaki, H. and Takahata, Y. (1990) Demography and Reproductive Profiles. In: Nishida, T. (Ed.) The Chimpanzees of the Mahale Mountains: Sexual and life history strategies. University of Tokyo Press: Tokyo, pp. 63–97.

Norconk, M.A. (2006) Long-term Study of Group Dynamics and Female Reproduction in Venezuelan *Pithecia pithecia*. International Journal of Primatology, 27(3), 653–674. Available from: https://doi.org/10.1007/s10764-006-9030-7

Norconk, M.A., Rosenberger, A.L. and Garber, P.A. (Eds.) (1996) Adaptive Radiations of Neotropical Primates. Springer Science & Business Media.

Okamoto, K., Matsumura, S. and Watanabe, K. (2000) Life history and demography of wild moor macaques (*Macaca maurus*): Summary of ten years of observations. American Journal of Primatology, 52(1), 1–11. Available from: https://doi.org/10.1002/1098-2345(200009)52:1<1::AID-AJP1>3.0.CO;2-F

Omar, A. and Vos, A. de (1971) The annual reproductive cycle of an African monkey (*Cercopithecus mitis kolbi* Neuman). Folia Primatologica; International Journal of Primatology, 16(3), 206–215. Available from: https://doi.org/10.1159/000155402

Ostner, J. and Kappeler, P.M. (2004) Male life history and the unusual adult sex ratios of redfronted lemur, *Eulemur fulvus rufus*, groups. Animal Behaviour, 67(2), 249–259. Available from: https://doi.org/10.1016/j.anbehav.2003.05.012

Pereira, M.E. (1991) Asynchrony within estrous synchrony among ringtailed lemurs (Primates: Lemuridae). Physiology & behavior, 49(1), 47–52. Available from: https://doi.org/10.1016/0031-9384(91)90228-G

Petersdorf, M., Weyher, A.H., Kamilar, J.M., Dubuc, C. and Higham, J.P. (2019) Sexual selection in the Kinda baboon. Journal of Human Evolution, 135, 102635. Available from: https://doi.org/10.1016/j.jhevol.2019.06.006

Phiapalath, P., Borries, C. and Suwanwaree, P. (2011) Seasonality of group size, feeding, and breeding in wild red-shanked douc langurs (Lao PDR). American Journal of Primatology, 73(11), 1134–1144. Available from: https://doi.org/10.1002/ajp.20980

Pullen, S.L., Bearder, S.K. and Dixson, A.F. (2000) Preliminary observations on sexual behavior and the mating system in free-ranging lesser galagos (*Galago moholi*). American Journal of Primatology, 51(1), 79–88. Available from: https://doi.org/10.1002/(SICI)1098-2345(200005)51:1<79::AID-AJP6>3.0.CO;2-B

Qi, X.-G., Li, B.-G. and Ji, W.-H. (2008) Reproductive parameters of wild female *Rhinopithecus roxellana*. American Journal of Primatology, 70(4), 311–319. Available from: https://doi.org/10.1002/ajp.20480

Radhakrishna, S. and Singh, M. (2004) Reproductive biology of the slender loris (*Loris lydekkerianus lydekkerianus*). Folia Primatologica; International Journal of Primatology, 75(1), 1–13. Available from: https://doi.org/10.1159/000073424

Rahaman, H. and Parthasarathy, M.D. (1969) Studies on the social behaviour of bonnet monkeys. Primates; Journal of Primatology, 10(2), 149–162. Available from: https://doi.org/10.1007/bf01730980

Rhine, R.J., Wasser, S.K. and Norton, G.W. (1988) Eight-year study of social and ecological correlates of mortality among immature baboons of Mikumi National Park,

Tanzania. American Journal of Primatology, 16(3), 199–212. Available from: https://doi.org/10.1002/ajp.1350160303

Robinson, J.G. (1988) Demography and Group Structure in Wedgecapped Capuchin Monkeys, *Cebus Olivaceus*. Behaviour, 104(3-4), 202–232. Available from: https://doi.org/10.1163/156853988X00520

Rowell, T.E. (1966) Forest living baboons in Uganda. Journal of Zoology, 149(3), 344–364. Available from: https://doi.org/10.1111/j.1469-7998.1966.tb04054.x

Rudran, R. (1973) The reproductive cycles of two subspecies of purple-faced langurs (*Presbytic senex*) with relation to environmental factors. Folia Primatologica; International Journal of Primatology, 19(1), 41–60. Available from: https://doi.org/10.1159/000155517

Rudran, R. (1977) Socio ecology of the blue monkeys (*Cercopithecus mitis stuhlmanni*) of the Kibale Forest, Uganda. Smithsonian Contributions to Zoology, (249), 1–88. Available from: https://doi.org/10.5479/si.00810282.249

Rumiz, D.I. (1990) *Alouatta caraya*: Population density and demography in Northern Argentina. American Journal of Primatology, 21(4), 279–294. Available from: https://doi.org/10.1002/ajp.1350210404

Savini, T., Boesch, C. and Reichard, U.H. (2008) Home-range characteristics and the influence of seasonality on female reproduction in white-handed gibbons (*Hylobates lar*) at Khao Yai National Park, Thailand. American Journal of Physical Anthropology, 135(1), 1–12. Available from: https://doi.org/10.1002/ajpa.20578

Setz, E. and Gaspar, D. (1997) Scent-marking behaviour in free-ranging golden-faced saki monkeys, *Pithecia pithecia chrysocephala*: sex differences and context. Journal of Zoology. Available from: https://doi.org/10.1111/j.1469-7998.1997.tb04852.x

Sharma, A.K., Singh, M., Kaumanns, W., Krebs, E., Singh, M. and Kumar, M.A. et al. (2006) Birth Patterns in Wild and Captive Lion-Tailed Macaques (*Macaca silenus*). International Journal of Primatology, 27(5), 1429–1439. Available from: https://doi.org/10.1007/s10764-006-9077-5

Shil, J., Biswas, J. and Kumara, H.N. (2020) Influence of habitat conditions on group size, social organization, and birth pattern of golden langur (*Trachypithecus geei*). Primates; Journal of Primatology, 61(6), 797–806. Available from: https://doi.org/10.1007/s10329-020-00829-y

Solanki, G.S., Kumar, A. and Sharma, B.K. (2007) Reproductive Strategies of *Trachypithecus pileatus* in Arunachal Pradesh, India. International Journal of Primatology, 28(5), 1075–1083. Available from: https://doi.org/10.1007/s10764-007-9204-y

Stone, A.I. and Ruivo, L.V.P. (2020) Synchronization of weaning time with peak fruit availability in squirrel monkeys (*Saimiri collinsi*) living in Amazonian Brazil. American Journal of Primatology, 82(7), e23139. Available from: https://doi.org/10.1002/ajp.23139

Strier, K.B., Mendes, S.L. and Santos, R.R. (2001) Timing of births in sympatric brown howler monkeys (*Alouatta fusca clamitans*) and northern muriquis (*Brachyteles arachnoides hypoxanthus*). American Journal of Primatology, 55(2), 87–100. Available from: https://doi.org/10.1002/ajp.1042

Strier, K.B. and Ziegler, T.E. (1994) Insights into ovarian function in wild muriqui monkeys (*Brachyteles arachnoides*). American Journal of Primatology, 32(1), 31–40. Available from: https://doi.org/10.1002/ajp.1350320104

Struhsaker, T.T. (1977) Infanticide and social organization in the redtail monkey (*Cercopithecus ascanius schmidti*) in the Kibale Forest, Uganda. Zeitschrift Für

Tierpsychologie, 45(1), 75– 84. Available from: https://doi.org/10.1111/j.1439-0310.1977.tb01009.x

Tecot, S.R. (2010) It's All in the Timing: Birth Seasonality and Infant Survival in *Eulemur rubriventer*. International Journal of Primatology, 31(5), 715–735. Available from: https://doi.org/10.1007/s10764-010-9423-5

Tian, J.-D., Wang, Z.-L., Lu, J.-Q., Wang, B.-S. and Chen, J.-R. (2013) Reproductive parameters of female *Macaca mulatta tcheliensis* in the temperate forest of Mount Taihangshan, Jiyuan, China. American Journal of Primatology, 75(6), 605–612. Available from: https://doi.org/10.1002/ajp.22147

Tinsley Johnson, E., Snyder-Mackler, N., Lu, A., Bergman, T.J. and Beehner, J.C. (2018) Social and ecological drivers of reproductive seasonality in geladas. Behavioral Ecology: Official Journal of the International Society for Behavioral Ecology, 29(3), 574–588. Available from: https://doi.org/10.1093/beheco/ary008

Trébouet, F., Malaivijitnond, S., Reichard, U. H. (2021) Reproductive seasonality in wild northern pig-tailed macaques (*Macaca leonina*). Primates, 62(3), 491–505. Available from: https://doi.org/10.1007/s10329-021-00901-1

van Noordwijk, M.A. and van Schaik, C.P. (1999) The effects of dominance rank and group size on female lifetime reproductive success in wild long-tailed macaques, *Macaca fascicularis*. Primates; Journal of Primatology, 40(1), 105–130. Available from: https://doi.org/10.1007/BF02557705

Vanthomme, H. (2010) L'exploitation durable de la faune dans un village forestier de la République Centrafricaine: une approche interdisciplinaire. MNHN.

Volampeno, M.S.N., Masters, J.C. and Downs, C.T. (2011) Life history traits, maternal behavior and infant development of blue-eyed black lemurs (*Eulemur flavifrons*).

American Journal of Primatology, 73(5), 474–484. Available from: https://doi.org/10.1002/ajp.20925

Watsa, M. (2013) Growing up tamarin: morphology, reproduction, and population demography of sympatric free-ranging *Saguinus fuscicollis* and *S. imperator*. Washington University in St. Louis.

Winkler, P., Loch, H. and Vogel, C. (1984) Life history of Hanuman langurs (*Presbytis entellus*). Reproductive parameters, infant mortality, and troop development. Folia Primatologica; International Journal of Primatology, 43(1), 1–23. Available from: https://doi.org/10.1159/000156167

Wu, A., Luo, Y., Wang, S., Chen, Z. and Wang, B. (2006) Preliminary study on breeding periodicity of wild francois' langurs (*Trachypithecus francoisi francoisi*) in Mayanghe Nature Reserve, Guizhou. Acta Theriologica Sinica, 26(3), 303.

Wu, H. and Lin, Y. (1992) Life history variables of wild troops of formosan macaques (*Macaca cyclopis*) in Kenting, Taiwan. Primates, 33(1), 85–97. Available from: https://doi.org/10.1007/BF02382764

Xiang, Z.-F. and Sayers, K. (2009) Seasonality of mating and birth in wild black-and-white snub-nosed monkeys (*Rhinopithecus bieti*) at Xiaochangdu, Tibet. Primates; Journal of Primatology, 50(1), 50–55. Available from: https://doi.org/10.1007/s10329-008-0111-4

Zhao, Q. and Deng, Z. (1988) *Macaca thibetana* at Mt. Emei, China: II. Birth seasonality. American Journal of Primatology, 16(3), 261–268. Available from: https://doi.org/10.1002/ajp.1350160307
